# Similar Dynamic Frontal Cortex Representations of Auditory Stimuli Cueing Opposite Actions and Rewards

**DOI:** 10.1101/2025.06.22.660924

**Authors:** Pingbo Yin, Susanne Radtke-Schuller, Jonathan B. Fritz, Shihab A. Shamma

## Abstract

Frontal Cortex (FC) plays a pivotal role in controlling actions and their dynamics in response to incoming sensory stimuli. We explored FC representations of the same stimuli when signifying diametrically opposite behavioral meanings depending on task context. Two groups of ferrets performed Go-NoGo auditory categorization tasks with opposite contingencies and rewards, and varied stimuli. Remarkably, despite the opposite stimulus-action associations, single-unit responses were similar across all tasks, being more sustained and stronger to (Target) sounds signaling a change in action, than to (Reference) sounds indicating maintenance of ongoing actions, especially during task engagement. Three major dynamic response profiles were extracted from the overall responses, and their combination defined separate neuronal clusters that exhibited different roles in relation to task events. Decoding based on the temporal structure of the population responses revealed distinct decoders that were aligned to different task events. Furthermore, the β-band power, extracted from the FC local field potentials, was similarly and strongly modulated during Target stimuli in all tasks despite opposite behavioral actions. Based on these findings, we propose a model of pathway-specific functional projections from the tripartite FC neuronal clusters to the basal ganglia that is consistent with previous evidence for the conjoint roles of the FC and striatum in adaptive motor control.

## INTRODUCTION

The frontal cortex (FC) has been implicated in many of the cognitive and executive control functions required for goal directed behavior (Komura et al., 2001; Bruni et al., 2015; Friedman & Robbins 2021; Duan et al., 2021), including reward-based decision-making (Coley et al., 2021; Liu et al 2021), response inhibition (Schiller et al., 2014; Li et al., 2020), working memory (O’Reilly 2006; Miller et al., 2018; Wilhelm et al., 2023), attentional control (Zikopoulos & Barbas 2007; Gregorlou et al., 2014), and top-down adaptive modulation of sensory filters to optimize behavior after task reversals (Banerjee et al., 2020). In the auditory system, cortical neurons can rapidly and dynamically adapt their spectrotemporal selectivity reflecting changing stimulus context and task conditions (Fritz et al., 2003, 2005; Yin et al 2014; Elgueda et al, 2019). This task-related receptive field plasticity may be shaped by changing functional connectivity between frontal cortex and auditory cortex (Fritz et al., 2010; Sheikhattar et al. 2018; Yin et al 2020). Context can transform the behavioral meaning of incoming stimuli and even cause the same sound to mean two opposite things in different circumstances. For example, a referee’s whistle may mean “start play!” or “stop play!” depending on the context.

In this study, we explored the role of the FC in this adaptive decision-making process by employing the same sounds to signify diametrically opposite meanings depending on task context and reward valence. In one type of task, upon hearing a Target sound the animal initiated licking to obtain a water reward (positive reward; **P-paradigm**). In the other task, it learned to *stop* licking for water when presented with the same Target stimulus in order to avoid a mild shock (negative reward; **N-paradigm**). In an earlier study (David et al., 2012) we found that such different task reward structures and stimulus-action contingencies, induced two strikingly distinct forms of receptive field plasticity in primary auditory cortex (A1). In light of the strong top-down projections from FC to auditory cortex (AC) influencing dynamic sensory filters (Caras and Sanes, 2017; Winkowski et al., 2018; Bimbard et al., 2018; Schneider et al., 2018; Mitteladt & Kanold 2023; Macedo-Lima et al. 2024), we wondered whether the differential receptive plasticity was driven by distinct FC representations of the two opposite paradigms.

Therefore, in the experiments reported here, we examined how the FC encodes the execution and inhibition of these opposite task behaviors. To do so, we trained two groups of ferrets on two auditory categorical Go-NoGo tasks, requiring each group to discriminate non-compact sound categories (Yin et al. 2016, 2020). Task stimuli varied along two acoustic feature dimensions: spectral frequency (tones for the ***TN-task***) or *temporal* modulation rate (amplitude modulated white noise for the ***AM-task***). While all ferrets learned both categorization tasks, the two groups differed in their behavioral responses to the task-reward structures. As indicated above, in the **P-paradigm**, one group of ferrets learned to lick for water reward when Target stimuli were presented and refrained from licking to Reference stimuli. In contrast, the group that learned the **N-paradigm** performed the opposite behavior and refrained from licking for water when Target stimuli were presented but could lick freely to Reference sounds (**Figure 1A**).

**Figure 1.**
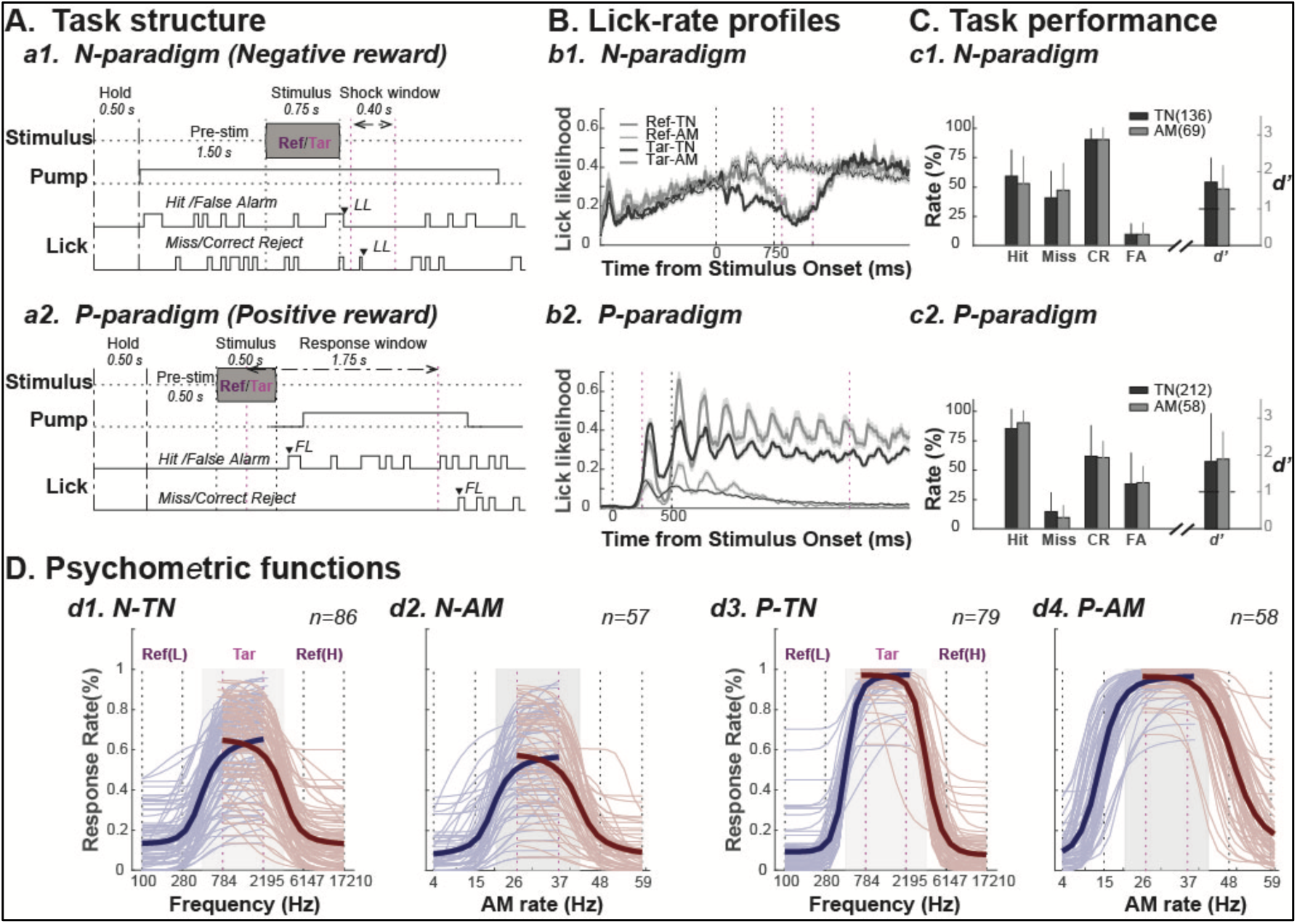
Behavioral Performance on Parallel Auditory Discrimination Paradigms with Negative or Positive Reward Structures. **A. Tasks structure**: Four task variants are illustrated in this panel. In all, trials were initiated when the animal refrained from licking the waterspout for 0.5 s. In the **N-paradigm (a1)**, after initial lick withholding, the water pump was turned on, and animals began licking for water. A task sound (either **Reference** or **Target**) was presented 1.5 s later. Animals learned to stop licking after Target sounds, but to continue licking to **Reference** sounds. Ferrets scored a **Hit** and avoided a mild electric shock if they withheld licking throughout a brief post-Target stimulus shock window (0.4 sec). However, if they licked during the shock window, they received a mild shock (**Miss**). By contrast, animals learned to continue licking after a **Reference** sound (**CR**, Correct Reject) and throughout the post-Reference stimulus period. If they stopped licking for the Reference, they received a 3-to-5 second timeout (**FA**, False Alarm) till the end of the trial. The last lick offset time (**LL**) before the end of the shock window was recorded and used to assess task performance. In the **P-paradigm (a2)**, a **Reference** or a **Target** sound was presented 0.5 s after a trial was initiated. After a **Reference** stimulus, animals learned to continue withholding licking (**CR**) from the spout. If they licked after a **Reference** stimulus, they received a 3-to-5 s timeout (**FA**) by the end of the trial. After a **Target** sound, the animals learned to initiate licking within the response window (**Hit**), triggering the water pump to turn on for 1.2-1.8 s (providing *∼0.2 ml* of water reward). The first lick onset time (**FL**) after a stimulus was recorded and used to assess task performance. **B. Lick-rate profiles:** Average licking profiles for all **Reference** and **Target** trials from all animals in each of the four task variants. The shaded area indicates standard errors. The vertical lines mark stimulus duration (dotted black) and response window (dashed magenta). **C. Task performance**: The bar plots show the average response rate for 4 different trial outcomes (**Hit**, **Miss**, **CR** and **FA**), and task performance (*d’*) over all behavioral sessions from all different task variants. Performance of individual animals is shown in Supplementary Figure 1A. **D. Psychometric Functions (PFs)**: PFs computed by fitting the likelihood of behavioral response at the borders between **Reference** and **Target** stimulus categories for each behavioral session (thin lines), with the thick lines being the average **PFs** across all *n* sessions during physiological recordings. The light grey shaded area marks the borders between the **Target** category (intermediate region) and each of the low (*blue lines*) and high (*red lines*) **Reference** category regions. The vertical dashed lines indicate the tone frequencies (or **AM** noise rates) used during training. The plots display **PFs** from two animals, one for the N-paradigm (**d1** and **d2**), and another for the P-paradigm (**d3** and **d4**). Both animals were trained on the same task stimulus sets of *tones* (**TN**-tasks) and *amplitude-modulated noise* (**AM**-tasks). The individual **PFs** from the remaining animals are available in Supplementary Figure 1B.

Extensive recordings during passive listening and active task performance allowed us to view in detail how task-salient stimulus features were represented in FC activity in the four variant behaviors comprising each of the two types of auditory categorization tasks and the two different reward-valence paradigms. The most surprising finding was a remarkable similarity of single-unit response patterns in all behavioral contexts despite opposite auditory-motor associations in the two paradigms. This result suggests a highly abstract level of representation in the FC. Another interesting result was the discovery of three distinct neuronal clusters with temporally-sequenced responses that aligned with the timing of behavioral responses of the animals in the two paradigms. A weighted composite of these three response clusters closely resembled β-band activity, derived from the accompanying local field potentials (LFPs), that have been shown to be a signature for either stopping ongoing actions, and/or switching and initiating new actions (Picazio et al., 2014; Shin et al., 2017; Jahanshahi et al., 2015; Wessel & Aron 2017; Wessel 2020; Diesburg et al., 2021; Horst et al., 2024; Polyakova et al 2024).

To integrate all these findings, we propose a testable model in light of the role of the FC-Basal ganglia loops (both cognitive and motor) in controlling and gating goal-directed behavior through three cortico-striatal pathways: the Hyperdirect, Direct and Indirect pathways (Alexander et al., 1986; Alexander & Crutcher, 1990; Mink 1996; Nelson & Kreitzer 2014; Jahanshahi et al., 2015; Baladron & Hamker 2020, 2024; Hernandez-Martin et al., 2023; Nambu et al., 2002, 2023; Redinbaugh & Saalmann, 2024). Many previous studies have explored the role of these descending cortico-striatal pathways in shaping behavior. Thus, according to classical basal ganglia Go-NoGo models, the Direct and Indirect pathways exert opposing control over movement (DeLong, 1990). However, more recent studies have shown that both pathways can be coactivated during intended movements (Cui et al., 2013) leading to cooperative, co-activation or complementary encoding (Bariselli et al., 2019; Varin et al., 2023), or task-dependent models (Bolkan et al., 2022). Our experimental results in the FC and proposed model provide new insights into delineating the contributions of these cortico-striatal pathways to adaptive movement.

## RESULTS

Five adult female ferrets (1-2 years old) were trained on auditory Go-NoGo tasks. After initial training, they were implanted with a headpost, habituated to head-fixation and retrained to behavioral criterion. In subsequent neurophysiological studies, we recorded single-unit activity from FC and AC in all five ferrets. Most of the data from the N-paradigm tasks have been previously analyzed to assess the categorical representations in primary and secondary AC, and FC (Yin et al 2020). In this study, we report FC data combined from both paradigms, but focused on the role of the FC in executing and controlling the animal actions in the two opposite behavioral tasks.

### Behavioral paradigms and stimuli

All ferrets were trained on both TN- and AM-tasks in which they discriminated two sound categories (Reference *versus* Target sounds) as detailed in **Table 1** in ***Methods***. Two animals learned to perform the N-Paradigm version of the task, while three others perform the P-Paradigm version (**Figure 1A**). Thus, there were four task variants in total (two auditory discrimination tasks in each of the N- and P-paradigms). Details of the task structures are depicted in **Figures 1A**, together with the definition of the various trial outcomes: **Hit**, **Miss**, Correct Rejection **(CR)**, False Alarm **(FA),** and Last Lick offset (**LL**), and how these outcomes are used to assess animal performance.

**Table 1:**
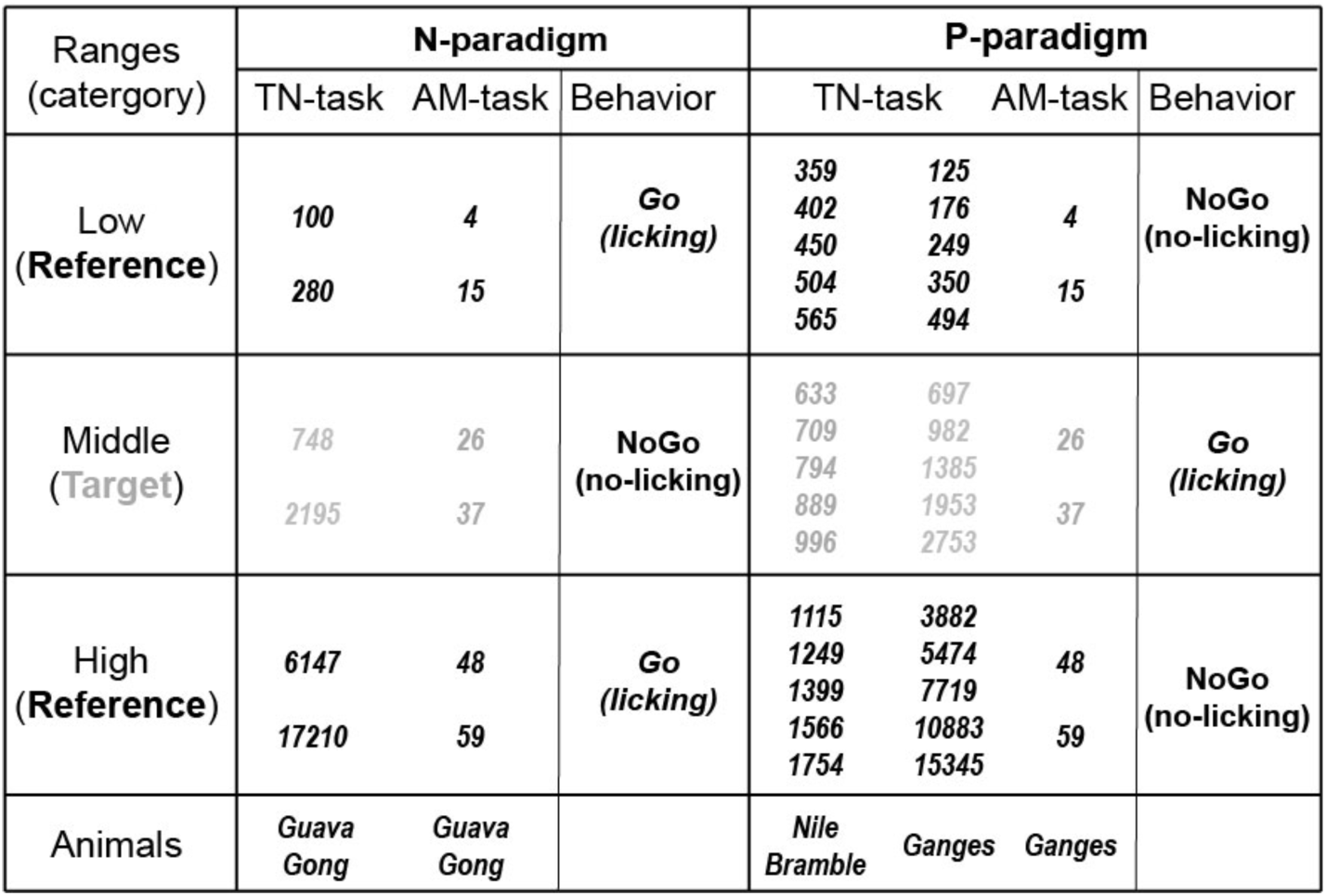
Training paradigms and stimulus sets.

All animals successfully learned to perform the tasks (with head-restraint in a recording stereotaxic frame), as evidenced by the group average behavioral performance across all recording sessions (**Figures 1B-C**). Performance of individual animals is provided in **Supplementary Figures 1A, 1B**. In the N-paradigm (**Figure 1B(b1)**), animals established a stable lick-rate after the water-pump was turned on at the beginning of the trial and maintained it during Reference sounds (*thin lines*). Lick-rate decreased significantly after Target sounds (*thick lines*). In the P-paradigm (**Figure 1B(b2)**), animals refrained from licking at the beginning of a trial, and exhibited a significantly higher lick-rate after Target sounds than to References. As seen in **Figure 1C**, significant performance was achieved in all four groups of paradigms and tasks, as assessed by ***d’*** (derived from ***Hit*** and ***FA*** rates). However, it is also evident that the type of performance errors reflected strongly the impulse of the animals to drink in the context of the two paradigms. Thus, most errors in the N-paradigm were ***Miss****’s* (**Figure 1C(c1)**) arising from animals failing to stop licking during the shock window after a Target sound. In contrast, in the P-paradigm (**Figure 1C(c2)**), most errors were ***FA****’s* as the animals failed to withhold licking after Reference sounds. To further characterize behavioral performance, we computed Psychometric Functions (PFs) by fitting the likelihood of behavioral response in relation to the stimulus metric with a 4-parameter logistic function (see **Methods***: Behavioral Assessment and Psychometric Function Fitting)*. Using these logistic functions, we could estimate the likelihood of a response error attributed to pure guessing (Guess Rate (***λ***) estimation) or to inattention or motor errors (Lapse Rate (***γ***) estimation). The animals likely developed distinct behavioral strategies in the two paradigms to maximize their water gain. For example, as shown in **Figure 1D**, N-paradigm animals (***d1, d2***) tended to downward-shift their PFs, choosing a *conservative* strategy or disengagement; while P-paradigm animals (***d3, d4***) tended to upward-shift the PFs (more evident in the AM-task and difficult potions of the TN-task; see **Supplementary Figure 1B**), choosing a *liberal* strategy or improved vigilance (see details in **Supplementary Figure 1C**). The different behavioral strategies were also evident by different single-unit activity driven by the error trials from the two paradigms (**Supplementary Figure 2**), and also by the dynamics of the choice probability which confirmed a significant behavioral modulation of the single-unit activity only during the P-paradigm (**Supplementary Figure 4**).

### Response types across different behavioral conditions and stimuli

In total we recorded 1480 neurons, with 601 neurons in the two animals performing the N-paradigm, and 879 neurons in the three animals trained on the P-paradigm (**Table 2**). Majority of recorded units were anatomically confirmed to be located in the dorsolateral FC (**dlFC**) regions ranging from the dorsal Prefrontal Cortex (**dPFC)** to the rostral part of Anterior-Sigmoid-Gyrus/Premotor-Cortex (**ASG/PMC**) (details in **Supplementary Figure 3A**) based on the ferret brain atlas (***Radtke-Schuller, 2018***). Single-units displayed a substantial diversity of responses to Target and Reference stimuli in all behavioral tasks as illustrated in **Figure 2** where they are depicted both as spike-rasters and post-stimulus time histograms (PSTHs). Responses displayed a striking enhancement for the active (task-engaged) condition, compared to the quiescent passive (task-disengaged) listening condition, irrespective of task reward paradigm or stimulus type (TN or AM). This enhanced response in the active state was most apparent in the Target responses but was also seen more modestly in the responses to Reference stimuli (**Figure 2b and Supplementary Figure 3B(a))**. While some neurons showed weak transient (**Figure 2a**) or sustained responses during the passive state (**Supplementary Figure 3B(a)),** the activity of many neurons was behaviorally gated – with virtually no responses to either Reference or Target stimuli in the passive condition, becoming vigorous only during task-engagement (**Figure 2b-f**). These findings are consistent with earlier results from FC (Fritz et al., 2010; Yin et al., 2020).

**Figure 2.**
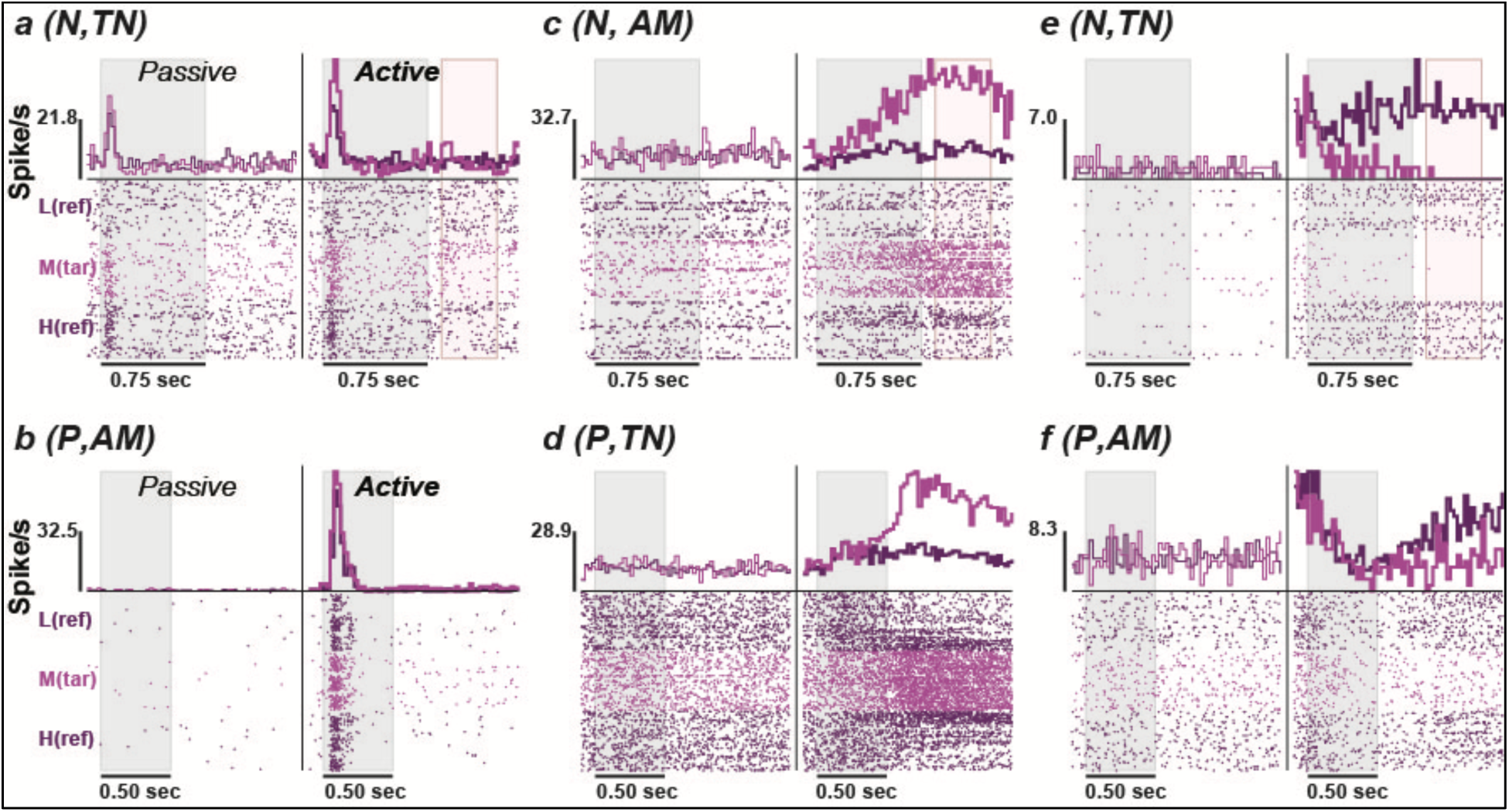
Diversity of single-unit responses in FC during performance of different tasks. Six representative examples of single-unit neuronal responses are shown, spanning both stimulus types (**TN, AM** noise) and the two reward N- and P-paradigms (**N, P**). In each of the 6 panels, the **Target** (**Reference**) responses in *pink* (*purple*) are shown in the passive (left-half of each panel) and active (right-half of panels) states. Overall, responses to **Targets** were stronger than to **References**, and greater in the active than in passive states. Five of the cells **(b-f)** exhibited behaviorally gated responses, being responsive only in the active task condition. Two cells (**a, b**) exhibited phasic responses to tones (panel **a**) or to AM noise (panel **b**). Both responded more vigorously to **Target** than **Reference**, and more strongly during the active than passive states. Two cells (**c, d**) showed gradual buildup of a sustained excitatory response to AM (panel **c**) and Tone (panel **d**) stimuli. A different pattern was seen in cells (**e, f**) that exhibited an increase in *baseline* firing rate during active task engagement while responding to task stimuli with sustained suppression both to tones (panel **e**) and AM noise (panel **f**) stimuli. The suppression was deeper during **Targets** than in **References**, and in the active compared to passive states.

**Table 2:**
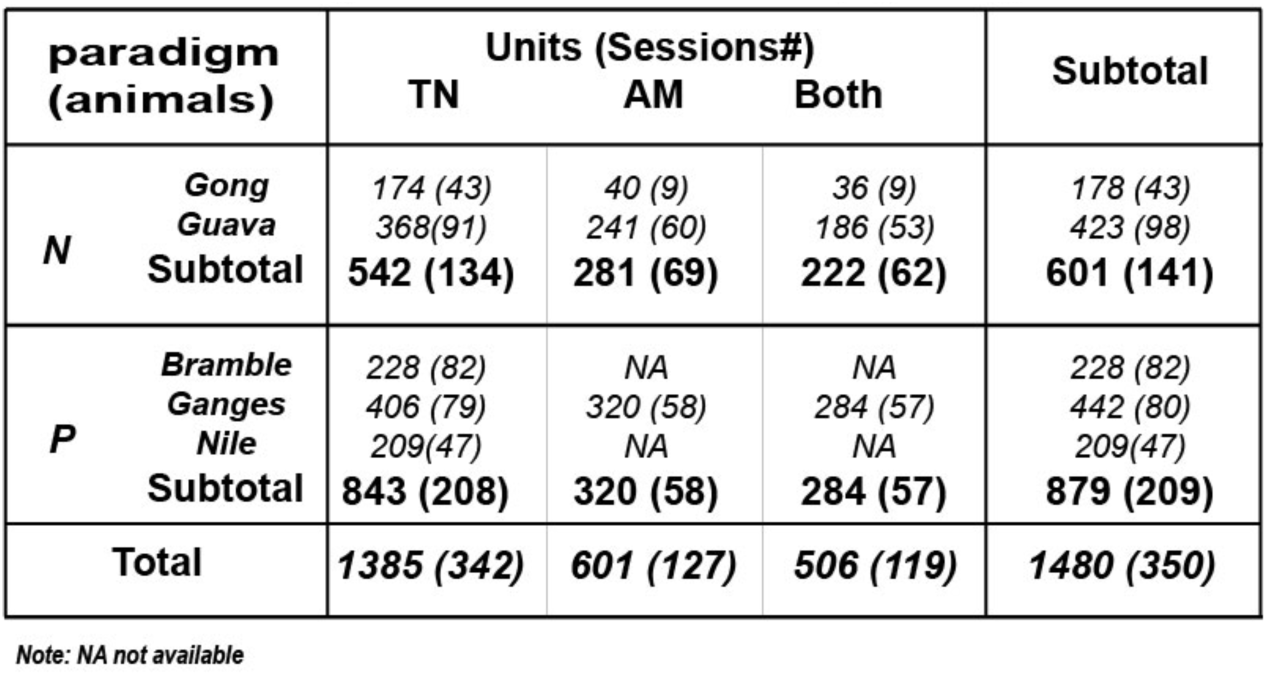
Summary of the isolated units and physiological sessions.

| paradigm<br>(animals) |  | Units (Sessions#) |  |  | Subtotal |
| --- | --- | --- | --- | --- | --- |
|  |  | TN | AM | Both |  |
| N | Gong | 174 (43) | 40 (9) | 36 (9) | 178 (43) |
|  | Guava | 368(91) | 241 (60) | 186 (53) | 423 (98) |
|  | Subtotal | 542 (134) | 281 (69) | 222 (62) | 601 (141) |
| P | Bramble | 228 (82) | NA | NA | 228 (82) |
|  | Ganges | 406 (79) | 320 (58) | 284 (57) | 442 (80) |
|  | Nile | 209(47) | NA | NA | 209(47) |
|  | Subtotal | 843 (208) | 320 (58) | 284 (57) | 879 (209) |
| Total |  | 1385 (342) | 601 (127) | 506 (119) | 1480 (350) |

Note: NA not available

Response patterns varied widely across the neuronal population, ranging over three major types: *transient* (or phasic) (**Figure 2**–*left panels)*, *sustained excitatory* (**Figure 2**–*middle panels*), *sustained suppressed* responses (**Figure 2**–*right panels*) with some neurons displaying hybrid responses (**Examples in Supplementary Figure 3B(b,h)**). However, for any given neuron, its response type was generally similar in all conditions, although displaying some modulation in strength depending on behavioral context (passive/active) and stimulus type (Target/Reference). For example, the two cells shown in **Figure 2(a,b)**, from different animals and reward paradigms, exhibited phasic responses in both passive and active conditions, to Reference and Target, and to TN and AM stimuli. The two other common response patterns (sustained excitatory and sustained suppressed responses shown in **Figure 2(c-f)**), were most strongly modulated by Target stimuli in the active condition. An interesting feature observed in many cells’ responses (**Figure 2(e, f)**) was a significant increase in baseline responses during the active relative to the passive states. The possible interpretation of the changes of the baseline activity during active state are discussed and shown in **Supplementary Figure 6.** We shall provide detailed analyses of these three basic response types and their potential functional significance across the entire FC neuronal population.

### Averaged population PSTHs exhibited similar temporal profiles across tasks

FC neurons exhibited similar response dynamics for encoding of categorical information in the two paradigms and tasks (see **Supplementary Figure 4**). To gain a broad view of the overall FC response patterns, the population PSTHs were assembled from isolated single-units in each of task variants (**Figure 3**), grouped by Target (*pink*) and Reference (*purple*) sound categories. Three features of FC activity emerged when viewing the population PTSH responses in the various conditions in **Figure 3**, namely: (1) responses to all task stimuli in the passive state (*thin line*s) were relatively weak and predominantly transient; (2) During the active state (*thick lines*), Target responses (*pink*) became strongly enhanced and more sustained in character, while (3) Reference responses (*purple*) remained far more phasic than Target responses. Reference responses were also significantly weaker in most conditions (except in the P-AM task (**Figures 3D**) where they were comparable). The enhancement of responses in the active state is highlighted by displaying the contrast (or difference) between the Active and Passive states, shown in the third column of all panels in **Figure 3**. An additional feature also emerged, namely: (4) a slightly delayed suppressive response to both Reference and Target stimuli, a response feature that (as we discuss later) may potentially play an important functional role in switching of ongoing behavior.

**Figure 3.**
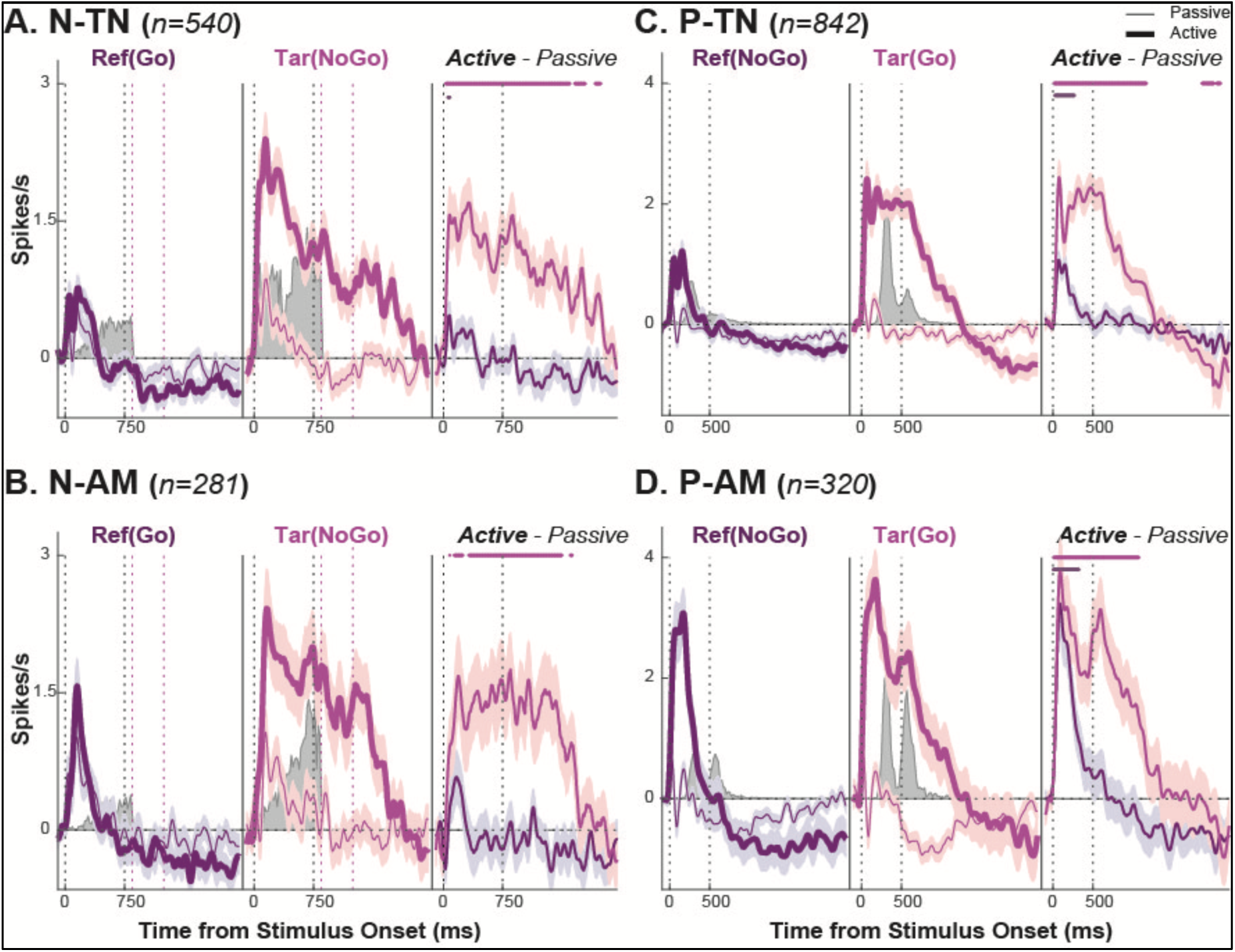
Distinct population response profiles evoked by Reference (Ref) and Target (Tar) stimuli in different task paradigms (N, P) and stimuli (TN, AM). The four panels are constructed from all FC neurons recorded in the **N**-(*panel*s *A, B*) and **P**-(*panels C, D*) paradigms, and both TN (*panels A, C*) and AM (*panels B, D*) stimulus types. The population PSTHs to Reference (*purple line, left subpanel*) and Target (*pink line, middle subpanel*) sound categories were corrected by the pre-stimulus baseline firing rate (measured in the 100 *ms* window prior to stimulus onset) in each experimental epoch. The PSTHs with *thin lines* indicate the response during the passive listening state, and the *bold lines* show responses during the active state (task engagement). The right subpanels of the plots depict the *net* changes in *Target* and *Reference* responses between the active *vs.* the passive states (i.e., Active *-* Passive). The lines at the top of each right subpanel indicate the significance of the changes (*by t-test, p<0.05 with Bonferroni correction for repeated measures*). The filled gray areas in each of the left and center subpanels display the distributions of the behavioral reaction times to Reference (*FA trials*) or Target (*Hit trials*) sounds (measured by first lick (*FL*) time after stimulus onset in the *P-paradigm*, or the last lick stop (*LL*) time in the *N-paradigm*, see Fig. 1A). The vertical black dotted lines indicate stimulus onset and offset time, and the shock window for the N-paradigm is indicated by the vertical *magenta* dotted lines. The shaded regions around each line are the standard errors at each moment. All PSTHs were smoothed with a *100 ms* Gaussian kernel (with *20 ms standard deviation*) sliding at 5 ms steps.

To summarize: Target responses in both reward paradigms exhibited stronger and more sustained responses compared to the phasic Reference responses. This similarity of FC responses across the two opposite paradigms is striking given the marked differences between the associated behavioral actions and strategies (see **Figure 1** and **Supplementary Figure 1**). Furthermore, since these similar FC responses preceded diverse behavioral reactions in the different task variants (as reflected by the overlaid distributions (*in gray*) of reaction times to Target (***Hits***) or Reference (***FA***) sounds in **Figure 3**), it suggests that the FC response does not precisely encode the motor details of the actual actions of licking or holding, but rather it likely broadly controls and gates these actions as we discuss at length later.

However, despite the overall similarity in the population responses across tasks and paradigms, there were a few features that distinguished the population PSTH responses between the various tasks which may be attributed to the details of the behavior or the diverse stimuli (**Figure 3**). For instance, one difference between the N- and P-paradigms (*left vs. right panels)* was the longer sustained Target responses of the P-paradigms (approximately 2 *vs* 1 sec), a difference that reflects specific differences between the durations of behavioral engagement (**Figure 1A**) following the Target stimuli in the two paradigms (i.e., in the N-paradigm the ferrets have to stop licking and to maintain this behavior beyond the shock window, whereas in the P-paradigm, the ferrets could initiate licking at any time within the response window). Another difference concerns the AM *vs.* TN categorization tasks (*lower vs. upper panels*). In the active state, onset responses to AM Reference stimuli were stronger than Reference responses to the TN stimuli in both the N- and P-paradigms, becoming comparable to (but still somewhat smaller than) the phasic portion of the Target response. One possible reason for the relatively strong Reference and Target phasic responses in the AM task is That the AM task is more difficult than TN task, therefore required relative more attention (and also longer reaction time) in performing the task, which might reflect in the phasic portion of the response (Yin et al, 2020).

### Principle components of the PSTH responses: three cell clusters

As described earlier (**Figure 2**), individual neurons exhibited a wide variety of response dynamics, especially to Target sounds during active task performance. Therefore, to further explore the nature of these response profiles, we performed Principal Component Analysis (PCA) over the data matrices formed by Time (the averaged PSTHs evoked by correct Reference/Target trials) x neurons (as detailed in **Methods** and **Supplementary Figure 5A-B**). In total, data from 4 task variants (N- and P-paradigms x 2 stimuli (TN & AM)) were used; the top 3 principle components (PCs) constituted a significant contribution to explaining the total variance across the neuron population (*>5% dashed line* in **Figure 4A**) and exhibited comparable response profiles (***Figure 4B***), in which the combination accounted for 67-84% of the total explained variance. We projected onto these 3 PCs all cell responses during the active task-engaged episodes, and K-means clustered the resulting coefficients (bottom panels of **Figure 4C**) into three groups (as detailed in **Methods** and **Supplementary Figure 5C-D**). The heat maps of the raster responses of the cells, as sorted by these clusters, are shown in **Figure 4C** (*top row of panels*), segregated among the four task variants. The population PSTH’s corresponding to each cluster are plotted in **Figure 4C** (*bottom row of panels*). These population partitions reveal three basic response patterns that were remarkably consistent across all tasks: (1) a first cluster of neurons that were mostly weighted by *profile-2* and/or *profile-3* (in **Figure 4B** & **Supplementary Figure 6A**), which displayed a rapid *transient* response pattern depicted by the green waveforms (***Trans***); (2) A 2^nd^ cluster of neurons that was dominated by the extracted *profile-1* with *positive* weights (**Supplementary Figure 6A**), exhibited a longer latency sustained excitatory response as shown by the red waveforms (***Sus(+)***); (3) a 3^rd^ cluster that was dominated by the extracted *profile-1* with *negative* weights (**Supplementary Figure 6A**) and had a medium latency sustained inhibitory response depicted by the blue curves (***Sus(-)***).

**Figure 4.**
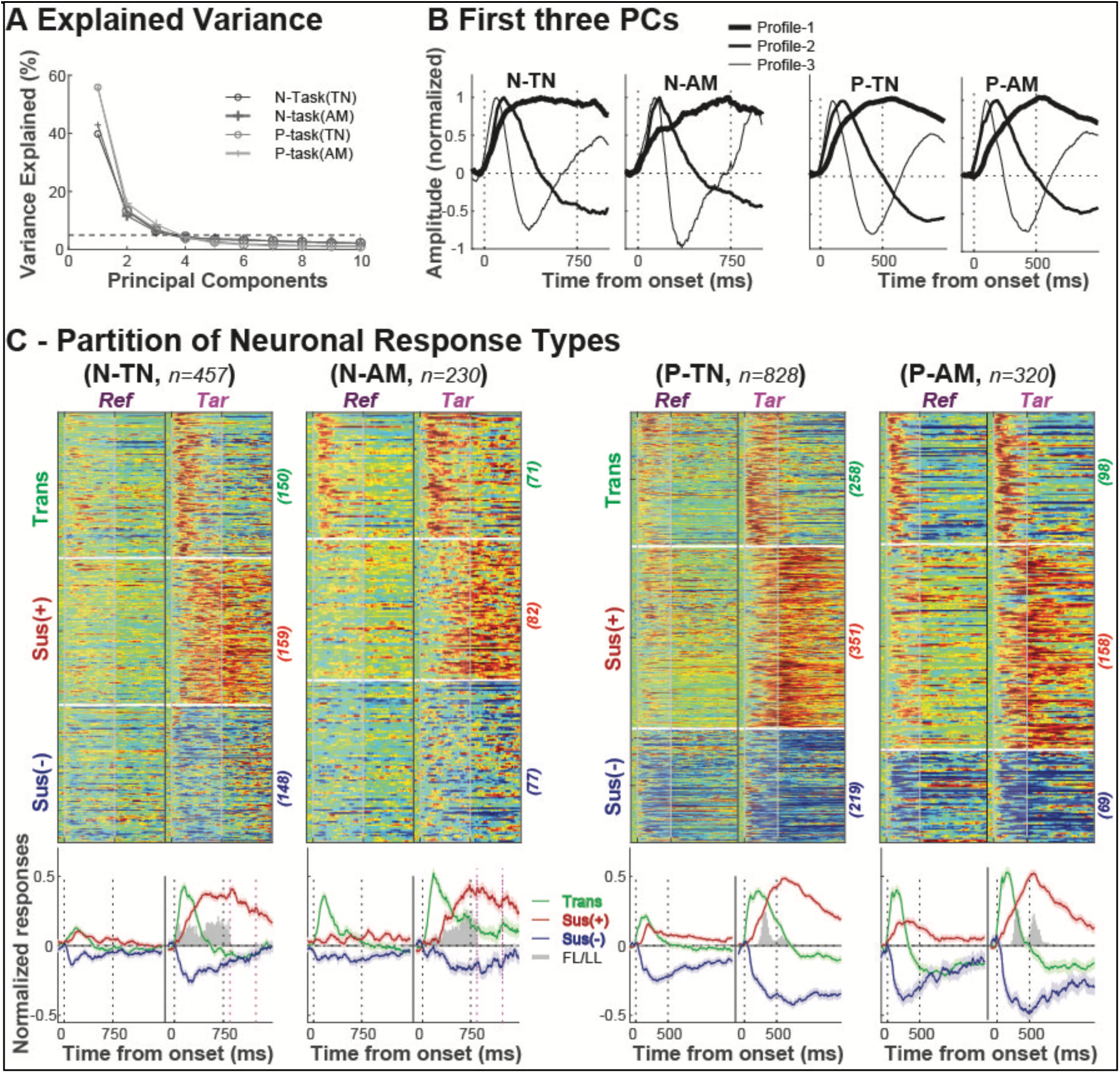
Principle component analysis and clustering of the FC neuronal responses. **(A)** PC analysis of responses in each of the four different task variants of Figure 3 yields 3 components, where each accounting over 5% of the variance (*horizontal dashed line*), and together account for an average of ∼75 % (*67.3, 68.2, 78.5 and 83.6*%) of the total profile variance among the analyzed time segments. **(B) The 3 largest PCs of each task variant** are similar across all task variants. **(C) Clustering all cell responses.** Computing the projection weights associated with the 3 PCs in each task yields three similar basic types of neuronal responses. Details of the clustering are given in **Methods** and **Supplemental** Figure 4. Each of the 4 panels shows the raster responses to the Reference (left) and Target (right) stimuli and are organized into 3 response types (clusters): Transient (***Trans***, *Top*); Sustained excitatory (***Sus(+)***, *middle*); Sustained inhibitory (***Sus(-)***, *bottom*). The overall average Reference and Target PSTH responses from each of the 3 cell clusters within a task are shown in the bottom row of panels. They are colored in green, red, and blue corresponding to the ***Trans****, **Sus(+)**,* and ***Sus(-)*** clusters. The filled gray areas display the distributions of behavioral reaction times to Target (*Hit trials*) for the N-paradigm (LL) and for P-paradigm (FL) as in Fig. 1A.

These three clusters emerged in both Target and Reference responses, varying only by their relative strength. In all cases, Target responses were significantly larger, and the combined three PC profiles gave rise to all the PSTH dynamics described earlier in **Figure 3**. The earliest transient response (*Trans* Cluster) is presumed to reflect in part the sensory and categorical information about the task stimuli (Yin et al. 2020), combined with information on the decision process, behavioral choice, and transformations to the selected action. The transient response is followed closely by a build-up of sustained responses both from the excitatory *Sus(+)* and inhibitory *Sus(-)* clusters which continue throughout the behavioral response of the animal to the Target stimulus. This sustained bidirectional (*positive/negative*) modulation of the response may reflect the initiation and maintenance of the selected action during the trial and is consistent with earlier findings (Figure 3 in Fritz et al., 2010).

Interestingly, neurons in both the *Trans* and *Sus(-)* clusters showed a tendency to increase their baseline firing rates (measured during a 100 ms pre-stimulus silence period) as the animal switched into the *active* task context from *passive* listening (see **Supplementary Figure 6B**). We reemphasize that the composition and dynamics of all task Reference and Target responses were similar, differing primarily by the relatively stronger Target responses in each of the three Clusters.

### Decoding reference and target stimuli based on population activity patterns

FC population activity integrated diverse neural signals that likely played different functional roles during task performance, such as categorizing the stimuli (Reference vs Target), decision making (**Go** vs **NoGo**) and execution/maintenance of selected actions (licking vs restraining licking). These multiplexed neural signals may have distinct temporal structures that can be disentangled and observed over the time course of the behavior (***Jagadisan & Gandhi, 2022***). To explore these dynamics in the Target and Reference population responses, we trained a decoder to discriminate between the two stimulus categories (details in **Methods**), by constructing first a pseudo-population 3D data matrix from segregated responses of the Reference and Target stimuli (**Figure 5A**). Each slice of the matrix included 12 population trials, with one randomly-selected correct trial from each neuron. To minimize overtraining effects, trials from each neuron were only selected once in each population draw. The resulting pseudo-population data matrix was then normalized by its Euclidean norm to form a unitary vector in multi-dimensional neuronal space at each time instant during a trial.

**Figure 5.**
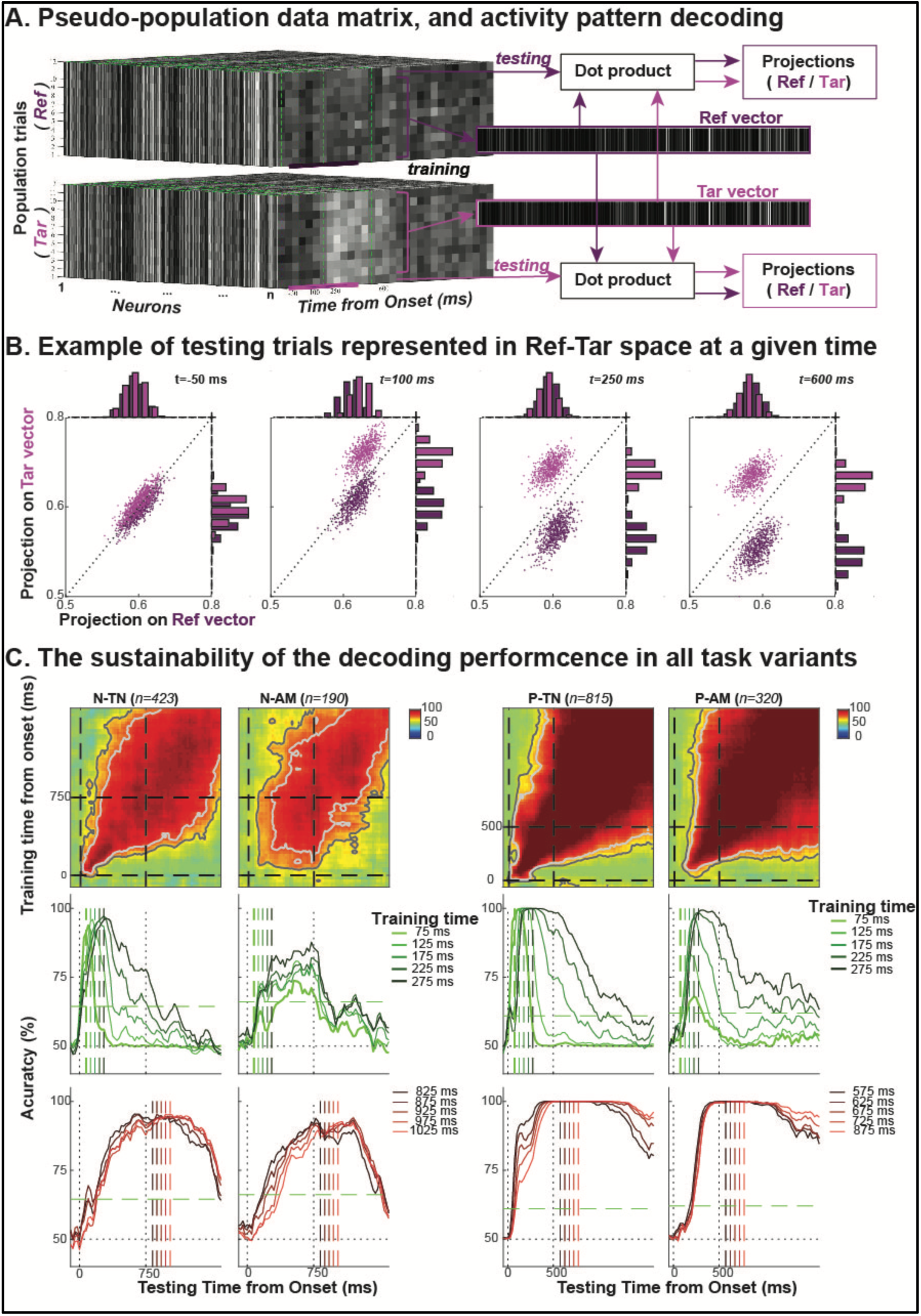
Decoding Reference and Target stimuli from population responses. **(A)** Training the Reference and Target vectors to decode the responses from each population trial. Details are provided in the text and in **Methods. (B) Scatter plot of response projections onto the Target and Reference decoders** (y- and x-axis, respectively). They are computed at different times throughout the trial as indicated in the different panels. Initially, responses are indistinguishable (pink/purple projections are mixed). However, they become segregated (decodable) within 100 ms from stimulus onset. **(C) Decoding dynamics are similar across task variants.** Each of the 4 columns of panels display the persistence of the response patterns over time based on their Reference/Target decoding. In the first row of panels, the heatmaps of the training *vs* testing times reveal similar patterns across the tasks. During the earliest phase, trained decoders change rapidly within 10’s of milliseconds (i.e., do not persist horizontally in time). This is also confirmed by the narrow cross-sectional patterns (middle row of panels) in the early training times (<100 ms) of the heatmaps (exhibiting narrow red regions around midlines). The maps exhibit significantly wider red regions at later times, indicating that trained decoders persist for hundreds of milliseconds. This is seen at times >150 ms (middle row of panels) and also in all last row of panels which display the cross-sections of the heatmaps at late training times. Further detailed interpretations of all these plots are available in the text and in **Methods.**

The flow chart extending to the right in **Figure 5A** illustrates the details of the decoding method with the population activation pattern using the “*leave-one-out*” cross-validation procedure (details in **Methods**). For each testing trial, the projections on the “Target-Vector” compared to the “Reference-Vector” reflected the likelihood at any time-point of the trial responses having been generated by either of the two categories.

An example of the results from this approach is shown with data from the P-paradigm, with a total of 1200 (100 draws) population trials for each stimulus type (**Figure 5B**). The scatterplots (Target *vs.* Reference trials) computed at different points in time from stimulus onset (−50, 100, 250 and 600 ms) illustrate the gradual change in the population responses, becoming reliably different for Target and Reference stimuli within 100 ms from stimulus onset. This is quantitatively confirmed by the bar plots along the two axes representing the distributions of the projections.

More insights into the decoding performance can be gleaned from the results shown in **Figure 5C**, where the topmost row of heatmaps illustrate decoding performance in each of the 4 task variants with the colormap indicating decoder training and testing accuracy at each time point from stimulus onset (chance performance at 50% level is colored green). Overlaid contour lines indicate one or two standard deviations above chance (>50% level is colored in progressively darker shades of red). The panels in the middle and bottom rows of **Figure 5C** illustrate cross-sections of the testing performance heat maps (**Figure 5C**; *top panels*) at various training times. For instance, training *early* in the trial (from 75 to 275 ms after sound *onset*, as depicted by the dashed vertical lines) reveals already above-chance discrimination between Target and Reference stimuli in all tasks. But the trained decoders from this early period only persist for tens of milliseconds. By contrast, when training the decoder later in the trial (175 to 275 ms after sound *offset*) when the animal was already engaged in its contingent behavior (**Figure 5C**; *bottom panels*), reliable discrimination emerged from as early as ∼300 ms after stimulus onset and the decoder remained stable throughout the rest of the behavior for hundreds of milliseconds.

These findings therefore suggest that the categorical decisions based on the sensory input that distinguishes Reference from Target stimuli likely occurred early following the onset of the responses (**Figure 5C**; middle row panels), and that they were mediated by the earliest transient (phasic) responses (Cluster 1 cells in **Figure 4**) that are similarly evoked early after onset. This is consistent with previous studies (Yin et al., 2020; see **Supplementary Figure 4**). In contrast, the behavioral actions associated with the two stimulus categories became stably encoded later in the trial (**Figure 5C**), suggesting that the combination of sustained responses of Clusters 2 & 3 populations were responsible for encoding actions elicited by the Reference and Target stimuli. This evolution of information in the population responses during the trial, from encoding stimulus category to encoding behavioral action, was also observed in the FC β-band responses that we discuss next.

### The β-band response patterns in the LFPs of the FC

One of the key objectives of this study was to explore how FC responses encode diverse actions driven by the same sounds in different behavioral contexts. Specifically, how are the opposite contingencies in response to Target sounds (i.e., start-licking in P-paradigms *vs* stop-licking in N-paradigms) reflected in the Target responses? Previous studies of stopping and switching behavior have emphasized the importance of β-band rhythms as LFP signatures that correlate with behavior (Diesburg et al., 2021; Tatz et al., 2023; Horst et al. 2024). We explored the dynamics and strength of the β-band rhythms as derived from LFPs recorded simultaneously with the FC single-unit data in the context of our two paradigms. Panels (**a1, b1**) in **Figures 6A, B** illustrate the grand average of the induced power spectrogram from wavelet transforms of the LFPs evoked by Reference (*top-half)* and Target (*bottom-half*) stimuli across all recording sites from the N-paradigm (**Figure 6A**) and the P-paradigm (**Figure 6B**). The data from the TN and AM tasks for each paradigm are combined (details of the wavelet transform computations and normalization are available in **Methods**). The power spectrograms for each task variant are provided in **Supplementary Figure 6**, which shows a comparable picture between TN- and AM-task within each of the paradigms. In both figures, panels are organized from left-to-right to depict responses in the **pre**-task passive (i.e. before task-engagement), **durin**g (i.e. active task performance), and the **post**-task passive behavioral states. The integrated power of the β-band responses (15—29 Hz) is displayed below in panels (**a2,b2**) for both Target (*pink*) and Reference (*purple*) responses in each state.

**Figure 6.**
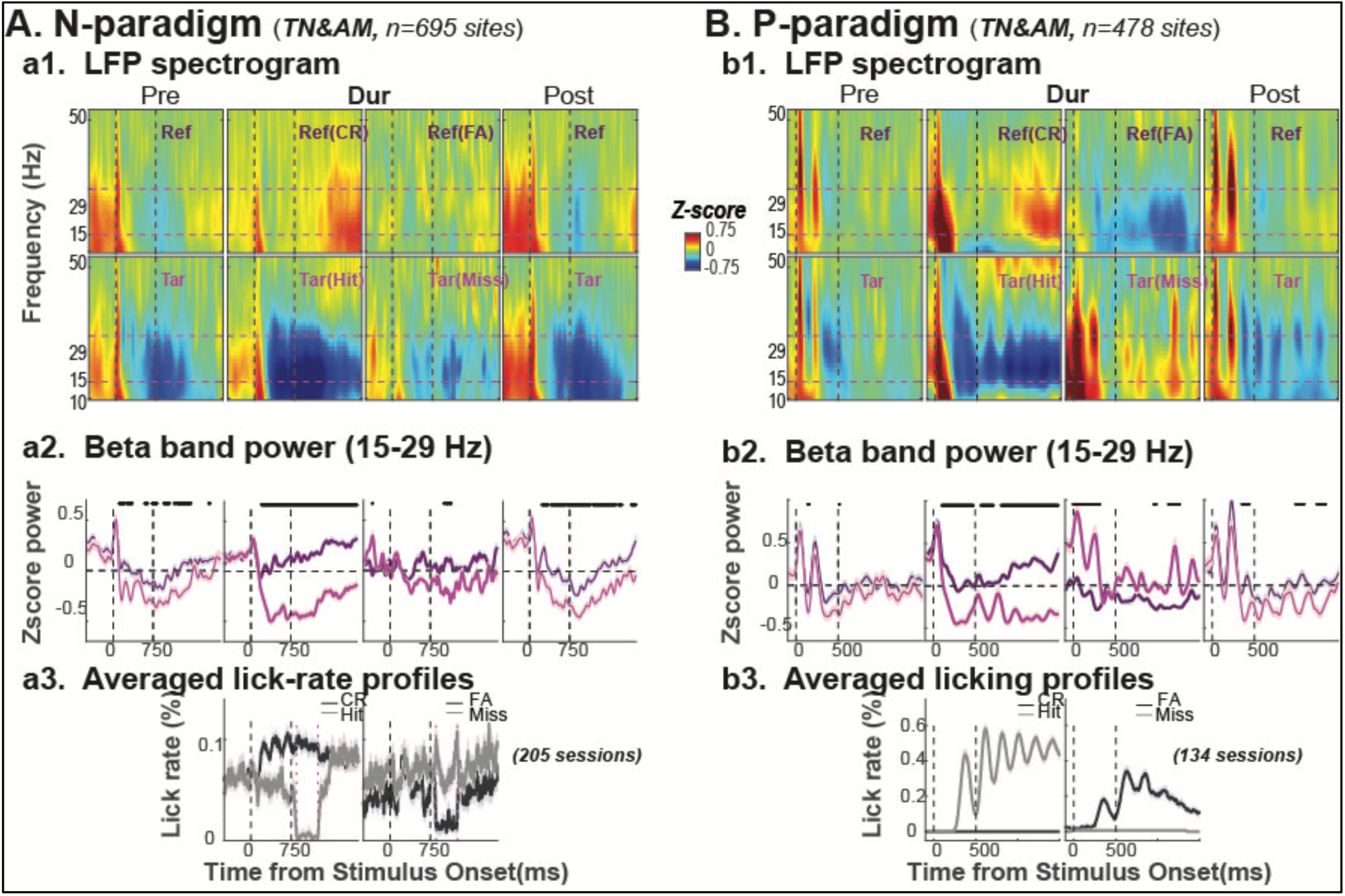
β-band (15-29 Hz) rhythms derived from LFP’s of FC responses. **(A)** β-band responses during N-paradigms. (***a1***) Each panel depicts the wavelet transforms of the LFPs near the β-band in the range marked by the horizontal dashed lines. Reference (‘**Ref’**) responses are shown in top row of panels from left to right during pre-passive (‘**Pre**’), active task engagement (‘**Dur**’), and post-passive (‘**Post**’). The second row of panels depicts the same information but for Target (‘**Tar**’) stimuli. The correct trials (*Correct Reject* (***CR***) and ***Hits***) and incorrect trials (***Miss*** and ***FA***) are displayed in separate panels during active task engagement. (***a2***) This row of panels contrasts the dynamics of the integrated β-band power from **Reference** and **Target** sounds. The black dots above the response profiles indicate significant differences between the induced β-band power of the two stimulus types (*by paired t-test, p<0.05 with Bonferroni correction for repeated measures*). (***a3***) Panels depict the lick-rate profiles for correct responses (***Hit*** and ***CR*** - left panel) and error responses (***Miss*** and ***FA*** - right panel) trials during active task engagement. **(B) β-band responses during P-paradigms.** The information depicted in all panels is analogous to that of the **A**-panels.

The β-band responses display a striking similarity in both behavioral paradigms during passive and active epochs as shown for the N-paradigm (**Figure 6: a1, a2**) and the P-Paradigm (**Figure 6: b1, b2**), consistent with the results from the single-unit data. For instance, the most prominent response dynamic in both paradigms occurs during the active Target responses (**Figure 6**: *middle panels* of **a1, a2**, **b1, b2**), where there was a brief, transient onset response to the stimulus followed by a deep depression of the β-band power that lasted throughout the behavioral response, usually recovering gradually back to baseline towards the end of the trial. Similar but much weaker dynamics were observed in the Reference responses, where the post-onset depression was shallower and recovered more rapidly. In the passive state (*left panels*), both Reference and Target response dynamics remained the same, albeit in weaker form. The response was seen most clearly in the post-task passive state where the memory of the active responses was clearly persistent. The β-band power depression during the active and post-passive responses are consistent with earlier single-unit results (e.g., ***Figure 6*** in Fritz et al., 2010).

When the animals committed behavioral errors such as ***FA*** (to References) or ***Miss*** (to Targets) during task performance, the β-band responses displayed different patterns as illustrated in the two middle (top and bottom) panels of **Figures 6A(a1,a2),6B(b1,b2)**. Broadly speaking, error responses in both the N- and P-paradigms are consistent with the animals’ cognitive state and their actions, i.e., Reference ***FA***’s resemble Target ***Hit****’s*, while Target ***Miss***’s resemble Reference ***CR***’s. Consequently, and as expected, the β-band power in Reference ***FA*** responses is *more* suppressed than ***CR***’*s*, whereas Target ***Miss*** responses become *less* suppressed than ***Hit’****s*. The amount of response modulations due to performance errors are far more salient in the P-than N-paradigms, a difference that likely reflects the divergent behavioral strategies in the two paradigms as discussed and illustrated previously in **Figure 1** and **Supplementary Figures 1,2**.

Finally, to emphasize the substantial dissociation between the FC β-band dynamics and licking, we show in **Figure 6A(a3), 6B(b3)** the lick-rate profiles during Target and Reference trials for both correct (*left panels*) and erroneous (*right panels*) trials in the N- and P-paradigms, respectively. Consider for example the elevated lick-rates following the Target in the P-paradigm (***Hit***-trace in *left panel of* **Figure 6(b3)**). By contrast, the lick-rate plunged to zero during the shock period of ***Hit*** trials in the N-paradigm (***Hit***-trace in left panel *of* **Figure 6(a3)**; 850-1250 ms interval between the two red dotted lines). Nevertheless, despite these opposite lick-rates in the two paradigms, the β-band suppression following the active Target (***Hit***) stimulus (**Figure 6 (**middle panels of **a1,a2** and **b1,b2)**) was similar in both the N- and P-paradigms. The same contrast between lick-rates and β-band responses applies to the Reference ***CR*** responses, where the absent *vs* sustained licking (***CR***-traces in left panels of **Figure 6(a3,b3)**) corresponds to a similar *shallow* depression in the Reference ***CR*** responses during the active state (**Figure 6(a1,a2** and **b1,b2)**).

## DISCUSSION

### Summary of the experimental results

This experimental study explored how behavior and its associated cognitive functions are encoded in ferrets’ frontal cortex as they categorized auditory stimuli in the context of Go-NoGo tasks (Yin et al. 2016, 2020). Ferrets learned to perform the tasks in one of two behavioral paradigms that were designed so that while the acoustic stimuli and categories were identical, the two paradigms were opposite in their reward valence and task structure. Adaptive decision-making in the frontal cortex is shaped by many factors including stimulus salience, the associated reward value, and valence of stimuli such as food (positive) or threat (negative). In the P-paradigm (positive reward) in this study, ferrets learned a “Go” response (start licking) in response to Target stimuli, whereas in the N-paradigm (negative reward), they learned a “NoGo” response (stop licking) to Target sounds to avoid a mild shock. Animals successfully learned to categorize both AM and TN stimuli in either of the two (Positive or Negative) reward paradigms but showed distinct behavioral strategies (**Figure 1** and **Supplement Figure 1**).

Although there was substantial diversity of single-unit responses during all stages of the tasks (**Figure 2** and **Supplement Figure 3)**, recordings nevertheless could be summarized succinctly by the average population PSTHs of **Figure 3**, where we observed substantially similar response features across all tasks despite the opposite contingencies and actions and behavioral strategies associated with the stimuli that induced them. Two general shared features were that all PSTHs exhibited larger responses during active than passive states, and greater responses to Target than to Reference stimuli (**Figures 2, 3**).

Response dynamics to task stimuli varied considerably especially across different behavioral conditions. For instance, during task performance, Reference responses were far more phasic (transient) compared to Target responses which displayed extended sustained responses over hundreds of milliseconds (**Figures 2, 3**). Another finding was that the PCA dimensionality reduction of all Target and Reference responses yielded three dominant PC’s that were similar across all task variants (**Figure 4A-B**) and explained about 65-80% of total response variance (**Figure 4A**). The population averaged responses could be shown to be composed of different weightings of these three PC profiles (**Figure 4C-D**). Based on these weights, the recorded FC responses could be clustered into three groups: *Trans, Sus(+), and Sus(-)* (**Figure 4C**). As a population, the early transient responses provided a reliable decoding of Target *vs* Reference stimuli beginning within 100ms after stimulus onset (**Figures 5B-C**). This gave way to a sustained and stable response over the subsequent hundreds of milliseconds (**Figure 5C**). We conjecture that the initial response phase in FC is related to stimulus categorization (**Supplemental Figure 6;** Yin et al., 2020). We further propose that this is immediately followed by a “decision” phase that induces the animal to stop ongoing action and switch to a different behavior. We suggest this represents an abstract, higher-level representation of stimulus-elicited change in goals and behavioral state (Koechlin et al. 2003; Gruber & McDonald, 2012; Vaidya and Badre, 2022; Baladron & Hamker 2020, 2024).

The β-band responses (extracted from the LFPs) also exhibited a remarkable uniformity across all tasks, with substantially greater response modulations to Targets than to References, and during active than passive states (**Figure 6A-B**). Active Target β-band responses consisted of a transient “positive” onset followed quickly by deep depression that was sustained throughout the subsequent behavioral episode. Passive Target responses exhibited a far shallower depression phase (**Figure 6A,B**), but with some persistence of the active target response during the subsequent post-passive periods, as previously seen in FC (Fritz et al. 2010).

Modulations of the β-band power in the FC and STN are seen as neurophysiological signatures of pausing, stopping or switching behavior in tasks with Go-NoGo or Pause-and-Switch behavior (Engel & Fries, 2010; Meyer & Bucci, 2016; Wessel & Aron, 2017; Wessel, 2020; Diesburg et al., 2021; Wessel & Anderson, 2024; Horst et al 2024). These two phases are related to the 2-stage model of response inhibition in Go-NoGo tasks (Schmidt et al. 2013; Schmidt & Berke, 2017; Diesburg et al.,, 2021; Hervault & Wessel, 2025; Wessel & Anderson, 2024), where a brief, initial burst of enhanced β-band activity is observed after Stop signals in a broad network including frontal cortex, Pre-SMA, M1, ventral thalamus and STN (subthalamic nucleus) (Wessel, 2020; Tatz, et al., 2023, Wessel and Anderson, 2024). This burst is postulated to represent an early “pause” process mediated by the Hyperdirect cortico-STN projection (Koketsu, et al. 2021), which provides a rapid mechanism for inhibiting or withholding task-related actions. A subsequent longer latency deep β-band depression (often termed a “Cancel” phase) is associated with voluntary *switching* from ongoing movements (Horst et al. 2024), which is what our animals do in response to Target stimuli.

### Functional relevance of target and reference responses

Planned movements activate an extensive brain network including frontal cortex, supplementary motor and motor cortices, basal ganglia and a set of cortico-striatal-thalamo-cortical pathways (Kravitz et al. 2010; Sippy et al., 2015). Although frontal cortex is known to play a key role in planning actions, stopping an ongoing action, or rapidly switching to a more optimal alternative action, we wondered what specific functional role the FC responses may have played in shaping the ferret behavioral responses in our study. Given the diversity of stimuli and behaviors across both tasks and reward paradigms, the uniformity of their associated responses in the FC is extraordinary and likely reflects a common functional thread of events with a shared meaning. To provide greater insight into the FC representation, we computed in **Figure 7A** the grand average of all response PSTHs (*leftmost panel*), of the three cluster responses (*Trans, Sus(+), and Sus(-)*) (*middle panels*), and of the β-band Target and Reference during the active states (*rightmost panel*). As expected, all closely recapitulate earlier response features, observed at the single unit, population, and LFP levels.

**Figure 7.**
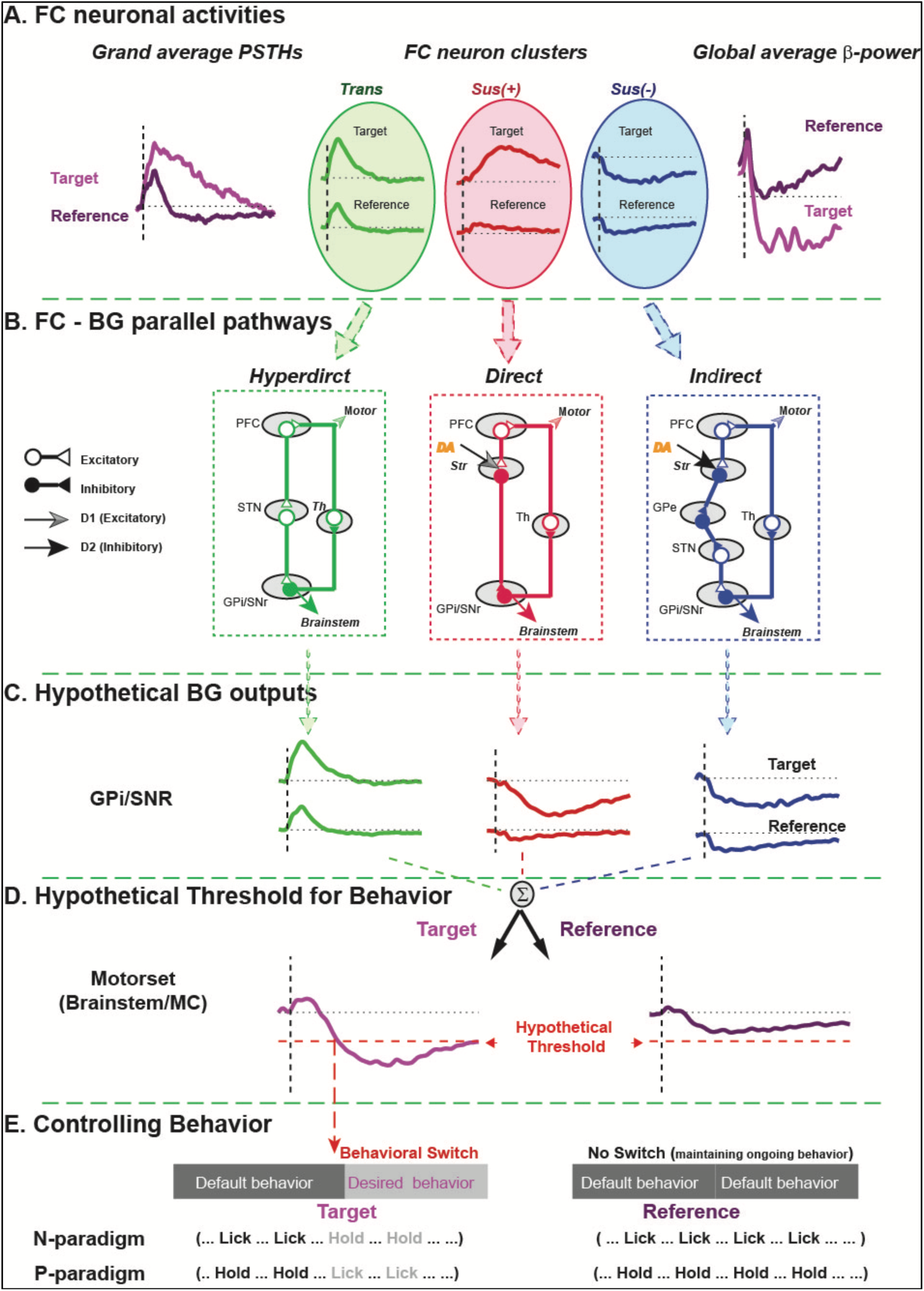
Hypothesized Functional Cortico-Striatal Projections from the Three FC Cell Clusters. **(A) Grand average FC neuronal responses**: Responses are averaged across all tasks and sources to generate grand average Target and Reference response PSTHs (left), three average cluster responses (*Trans, Sus(+),* and *Sus(-)*), and an average ゲ-band power. **(B) Simplified neuroanatomical layout of the three known major cortico-striatal pathways**: *Hyperdirect, Direct,* and *Indirect*. It is proposed that the three distinct FC cell clusters may project in parallel to these three pathways. Also shown are the Basal Ganglia (**BG**) schematic projections to the thalamus, and back to the FC to form the thalamo-cortical loops. The loop output also reaches to some motor areas (brainstem and motor cortex). Note that the dopamine (***DA***) signals from mid-brain structures (mainly the Substantia Nigra pas Compacta (***SNc***)) induce opposite effects on the direct and indirect striatal pathways through their characteristic ***DA*** receptors (excitatory ***D1*** vs inhibitory ***D2***). **(C) Conjecture that the BG output from the GPi/SNr** is the sum of *Trans, Inverted-Sus(+)*, and *Sus(-)* cluster responses originating from the FC. Target responses are larger than Reference responses but otherwise similar in form. **(D) Integrating the responses from all three BG pathways** at its output produces larger Target (left) and smaller Reference (right) responses that closely resemble the average β-band power responses depicted in the rightmost panel of Fig. 7A. **(E) Schematic of the functional role of Reference and Target** stimuli on task behavioral actions and performance in the N- and P-paradigms. The integrated activity of GPi/SNr induced by Target (but *not* Reference) stimuli exceeds a hypothetical negative threshold (the horizontal dashed lines and arrowheads depicted in Fig. 7D), and hence leads to behavioral switching from ongoing (default) behavior to task desired behavior by disinhibition of the downstream motor related regions (brainstem or alternately to motor cortex by way of the motor thalamus as depicted in Fig. 7B).

We next considered the possible common functional role that Target and Reference stimuli have in all tasks, particularly in the opposite contexts of the N- and P-paradigms. Previous studies have emphasized that FC encodes abstract, higher-level rules (Koechlin et al., 2003; Gruber & McDonald 2012; Vaidya & Badre, 2022; Baladron & Hamker, 2020, 2024). If the FC responses are interpreted as acting at a higher level of abstraction, then Target stimuli could serve to signal to the animal the need to switch (change) its ongoing behavior to an appropriate alternative. In contrast, Reference stimuli signal to the animal to maintain its current behavioral state. Therefore, in this view, Target responses in FC do not reflect details of the different motor-sets (licking *versus* non-licking) but rather mediate a behavioral change (switching actions), while Reference responses promote maintenance of the status quo and continuation of ongoing action (Engel and Fries, 2010). Our results show that Target sounds produce similarly sustained responses (at a single-cell, population level, and in the LFP β-band) whether the animal suddenly withholds licking (N-paradigm) or initiates licking (P-paradigm) (**Figures 3,6**). This abstraction of task structure is highlighted in the schematic of **Figure 7E** which translates the N- and P-paradigms to this shared and more abstract description (Koechlin et al. 2003; Vaidya and Badre, 2022).

Given the close interactions between the FC and Basal Ganglia (BG), one challenge we face in integrating our findings with the existing BG functional literature is that in most Go-NoGo, Stop signal reaction time (SSRT), and other Stop-Signal studies (Wessel & Anderson, 2024), there has been a literal interpretation of “Go” and “NoGo” choices as “start action” *versus* “stop action”. A central insight from our results is that FC responses, and by inference BG outputs (discussed next), do not necessarily reflect movement *per se*, but rather the abstract control of desired actions that may or may not include movement. Thus, withholding licking in the N-paradigm tasks (actively switching from licking to non-licking) is abstractly an action like any other, despite the apparent behavioral inaction (although there may well be more complex behavioral responses even during the NoGo Condition – for example the tongue may be actively retracted). Hence, we propose that the most general way to think about the change in behavior in Go-NoGo tasks is in terms of switching from one behavioral state to another.

### Possible contributions of the three FC clusters to the FC-Striatal projections

How do the three basic types of FC responses contribute to task behavior? It is known that frontal cortex projections to the basal ganglia are essential for movement control, and play a key role in action selection, initiation, and modulation of motor activity.

According to the classical basal ganglia motor model, two pathways - Direct and Indirect (**Figure 7B**) - exert opposing control over motor behavior. A third pathway - the STN (**Figure 7B**) - plays a pivotal role and receives inhibitory input from the *Indirect* pathway (via GPe) and also rapid excitatory input from the Hyperdirect pathway (cortex) and its output is excitatory. According to the classic model, intended “Go” movements are promoted via the *Direct* pathway (cortex -> striatum -> internal segment of the globus pallidus/substantia nigra (GPi/SNr) -> thalamus). By contrast, the Indirect pathway (cortex -> striatum - external segment of the globus pallidus (GPe) -> STN -> GPi -> thalamus), mediates suppression of competing motor programs, and conveys stop signals by a “No-Go” response (Nambu et al. 2002; Frank, 2005; Bariselli et al., 2018; Wessel, 2020).

We speculate that the interplay between these FC responses propagating through these three major cortico-striatal projections may constitute a fundamental mechanism underlying the way the FC reshapes or maintains actions in response to sensory stimuli as we elaborate next. **Figure 7B** depicts a simplified schematic of our proposed three major FC projections to the striatum. The basal ganglia in turn project back to the FC though the thalamus to form closed loops considered vital for the maintenance of ongoing actions. We conjecture that the three FC clusters may initiate and reflect the loop activity arising from the three corticostriatal pathways, in which the *Trans, Sus(+),* and *Sus(-)* clusters dominate the Hyperdirect, Direct and Indirect pathways, respectively. As shown in **Figure 7B**, when converging in the BG’s output nuclei (GPi/SNr), the Direct pathway *inverts* its FC *Sus(+)* inputs (via the inhibitory GABAergic striatal D1-MSNs), whereas the Indirect pathway *preserves* the polarity of FC *Sus(-)* inputs (through GABAergic D2-MSN neurons to the GPe). The third (Hyperdirect) *Trans* FC projection is an excitatory monosynaptic input to the STN, and the STN output is via excitatory glutamatergic neurons. Hence the Hyperdirect pathway is more rapid than the other two pathways. We propose that the three FC cluster activities converge on BG’s output nuclei (GPi/SNr), exhibiting a remarkably similar profile (**Figure 7C,D**) to the global average β-band power profiles (*rightmost panel* in **Figure 7A**).

The rationale for these hypothetical projections is the observed dynamic sequence of response features in our study that is broadly consistent with experimental evidence. The FC responses in our study suggest that initially, in the quiescent non-task-engaged state (passive listening) when there is minimal FC response to any acoustic stimuli, and hence no FC-striatal drive, the BG outputs of the GPi/SNr issue a sustained inhibitory output to the motor-set through the related motor regions (brainstem and/or motor cortex through motor thalamus) that maintains the current motor state of the animal. To initiate a different voluntary movement, we hypothesize that the FC output, sent down the different cortico-striatal pathways, must *reduce* the BG inhibitory output significantly, nominally below a threshold level (marked by the dashed red line in **Figure 7D**) so that the thalamo-cortical loop projection becomes excitatory. During task engagement, when strong FC responses to Target stimuli cause BG outputs to decrease below threshold, and release BG inhibition of the motor-set, we propose that it then becomes possible for the animal to switch or change a previous (default) action and initiate a different action. By contrast, since the FC responses to Reference stimuli are relatively weak, we suggest they would not reach threshold at the BG output (**Figure 7D**; right panel), thus maintaining the ongoing behavioral status quo, by continued inhibition of the motor-set.

The proposed model of interplay between the *Trans, Sus(-)* and *Sus(+)* responses is summarized in **Figure 7C**, leading to the composite in **Figure 7D**. In the active state, the earliest component of the Target response is the phasic *Trans*, projected through the Hyperdirect Cortico-STN pathway (**Figure 7B**), producing a phasic positive BG output (**Figure 7C,D**; left panels) that rapidly induces the animal to pause thalamic and cortical activity and interrupts ongoing actions (Schmidt & Berke, 2017; Koketsu et al. 2021). Next, to allow the animal to switch rapidly to a new behavior, we hypothesize that this initial Hyperdirect STN response is followed and supplanted (Lisman, 2014) by the addition of a fast-growing inhibitory *Sus(-)* component (**Figure 7B**) presumed to arrive soon after the Hyperdirect “pause” signal, through the (noninverting) Indirect pathway (**Figure 7C**; right panel). Thus, we propose that the net BG output (**Figure 7D**; left panel) begins to decrease below threshold, enabling a new behavior (motor-set) to commence. This in turn may generate additional thalamic excitatory activity that is propagated back to the FC through recurrent thalamocortical loops (Baladron & Hamker, 2020, 2024), inducing an additional gradual buildup of the FC excitatory *Sus(+)* response component (**Figure 7A**; middle panel). This increasing component projects back to the BG via the (inverting) Direct pathway (**Figure 7B**; middle panel) causing the composite BG output to decrease further below threshold (**Figure 7D**; left panel), thus reinforcing the new behavior. The proposition that the growth of *Sus(+)* underlies the buildup of either (opposite) Target actions is evidenced by its correlation with the buildup of the first-lick and last-lick distribution profiles, shown in **Figure 3**.

### Conclusions and future experimental validations

The proposed model (**Figure 7**) for the functional significance of the FC response clusters, their role in the control of actions during task engagement, and their segregation in specific fronto-striatal pathways is consistent with many aspects of the current literature. This schematic may in fact broadly underlie analogous tasks where a different action is initiated by a change from the *status quo* (or ongoing) action upon correctly perceiving a “Switch Cue” stimulus.

The involvement of the BG and thalamocortical loops in this process, and specifically the proposed tripartite projections of different FC clusters to the Striatum, and the hypothesized specificity of the cortico-striatal-thalamo-cortical loop projections are all highly speculative conjectures (Baladron & Hamker, 2020, 2024; Pasquereau & Turner, 2017, 2023). Other network mechanisms may play a role. For instance, recurrent FC activity can arise intrinsically within its microcircuits and ensembles even in untrained animals (Meyer et al., 2007; Nolan et al., 2025) and may therefore contribute to the nature of the FC responses in the cortico-striatal-thalamo-cortical loops that shape behavior. Further, our experiments have not demonstrated that the *same* neuronal clusters in FC are involved in mediating the responses to stimuli associated with both the P- and N-paradigms, since our recordings with the two opposite reward tasks were conducted in different animals. Hence, there is a need for further studies on FC responses in animals that have learned to switch between different paradigms in the same recording session (David et al., 2012; Bornhoft et al., 2025). This is important to resolve, especially because there is some evidence for regionally distinct areas of FC associated with appetitive or aversive stimuli (Monosov & Hikosaka, 2012).

In conclusion, our study suggests that the FC encodes a dynamic sequence of three major types of responses that encode stimulus identity, category, and behavior. Moreover, the similarity of FC responses in both the P- and N-paradigms, independent of behavior and associated reward or punishment, suggests that a key element of the FC responses in the Go-NoGo tasks in our study is whether to continue the status quo (ongoing) behavior or to switch to a new behavior. Our model seeks to integrate the FC responses with known cortical projection pathways to the Basal Ganglia. However, future experimental neurophysiological, neuroanatomical and optogenetic studies to test for the existence of segregated projections of FC clusters to the striatal inputs (Nonomura et al., 2018) will be essential to ascertain the validity of the conjectures underlying the proposed model. *This work was supported by grants from the National Institutes of Health (R01 DC005779)*

## AUTHOR CONTRIBUTIONS

P.Y. designed the experiments. P.Y. trained the animals and performed all neurophysiological recordings. P.Y. and S.A.S. analyzed data and prepared the figures describing the behavioral, neurophysiological results and principal component and decoding analysis. P.Y. made the iron deposits/dye injections. P.Y. and S.R.-S. analyzed the neuroanatomical results and prepared the neuroanatomical figures. P.Y., S.A.S., and J.B.F. discussed interpretation of the data, revised figures and wrote the manuscript.

## METHODS

All animal experimental procedures were conducted in accordance with the ***National Institutes of Health’s Guide for the Care and Use of Laboratory Animals*** and were approved by the Institutional Animal Care and Use Committee (***IACUC***) of the University of Maryland.

### Animals

Five adult female ferrets (*Mustela putorius*) obtained from Marshall Farms ***(North Rose, NY***) were used in the current study. Ferrets were housed in pairs or trios in animal facilities accredited by the Association for Assessment and Accreditation of Laboratory Animal Care (***AAALAC***) and were maintained on a 12-hr light-dark artificial light cycle. All ferrets began behavioral training between 1-2 years of age, with weights between 600-900g. During initial behavioral training and the following behavioral physiological recording period, animals were placed on a weekly water-control protocol, in which they received *ad libitum* water freely over weekends. On weekdays, they were brought to the laboratory for daily training/testing and returned to the animal facility after completion of their behavior session. While on training/testing during weekdays, the animals received water reward for correct trials during their behavior session. Additional water supplements were given after testing if an animal didn’t drink sufficiently during their daily behavior session. Animal health was carefully monitored daily by experimentalists and veterinary staff to avoid dehydration or weight loss (animals were maintained above 80% of *ad libitum* weight).

### Experimental apparatus

Behavioral training was conducted within a single-walled, sound attenuated chamber (***IAC, INC.***). Ferrets were trained in a custom-built transparent Lucite testing box (18 cm width X 34 cm depth X 20 cm height), placed inside the booth. A lick-sensitive waterspout (2.5 cm x 3.7 cm) stood in front of the testing box 12.5 cm above its base, positioned so that animals could easily lick from the waterspout to obtain water through a small opening in the front wall of the box. The waterspout was connected to a computer-controlled water pump (***MasterFlex L/S, Cole-Parmer***) for one training paradigm (the Negative Reward or ***N-paradigm***), or to a computer-controlled water dispenser (***Crist Instrument Co., Inc.***) for the other training paradigm (the Positive Reward or ***P-paradigm***). A custom-built interface box monitored the animal’s lick behavior, measured specifically by tongue contact with the waterspout, and converted the licks into a TTL digital signal. The TTL lick signal output was then fed back to the computer. A loudspeaker (***Manger, Germany***) was positioned 40 cm in front of the testing box for sound delivery during behavioral training, and the animal’s behavior was streamed with a video camera, displayed graphically trial-by-trial and continuously monitored on a computer screen. All acoustic stimuli used in the experiments were generated using MATLAB (***The MathWorks, Natick, MA***) at a 40 kHz sampling rate, and ramped with 5 ms rise-fall time. The sounds were converted at 16-bit resolution through a NI-DAQ card, then amplified and delivered to the loudspeaker. All behavioral training and the following neurophysiological recordings were controlled and monitored through a custom-built MATLAB GUI. All related trial events were recorded and stored in the computer for further analysis.

### Behavioral Paradigms

Two opposite variants of a Go/NoGo task with reversed stimulus-action behavioral contingencies were employed in the experiments in this study. Three ferrets (*Nile, Bramble, Ganges*) were trained on a three-range classification task using a positive-reinforcement paradigm (***P-paradigm***, for full details see ***Yin*** *et al*., 2016), in which animals were trained to lick a waterspout (***Go-behavior***) after a “safe” Target sound to obtain drops of water reward (*0.2∼0.4* cc), or to refrain from licking the waterspout (***NoGo-behavior***) following presentation of a “warning” Reference (non-Target) sound to avoid a 5-10 s timeout (*top panel in* ***Figure 1B***). The two other ferrets (*Gong, Guava*) were trained on the task using a conditioned avoidance paradigm (***N-paradigm***, for full details see ***Yin*** *et al*., 2020). In the ***N-paradigm***, animals began licking the waterspout for water when the water pump was tuned on (*at a flow rate of 0.6-1.2 ml/min*) at trial onset. They learned to stop licking the waterspout (***NoGo-behavior***) after the “warning” Target sound to avoid a mild electric shock either to the paw (or to the tail during physiological recording). However, following the presentation of a “safe” Reference (non-Target) sound, animals learned to continue licking for water (***Go-behavior***). Failure to lick to the Reference stimulus lead to a timeout after the trial (*top panel in* ***Figure 1A***). Hence, stimuli in the ‘***Target****’* sound category lead to ‘***Go-behavior***’ in animals trained with the ***P-paradigm***, and in contrast, lead to ‘***NoGo-behavior***’ in animals trained with the ***N-paradigm***. Stimuli in the ‘***Reference****’* sound category lead to ‘***NoGo-behavior***’ in animals trained with the ***P-paradigm***, and in contrast, lead to ‘***Go-behavior***’ in animals trained with the ***N-paradigm*** (see ***Table-1***). Thus, in the two animal training groups and associated two behavioral paradigms, the same stimulus categories (***Target*** *or **Reference***) were associated with opposite behavioral actions (***Go*** *or **NoGo*** licking responses).

In both training paradigms, the stimulus in each trial consisted of the presentation of a single sound (*0.5 s* duration in ***P-paradigm***, and *0.75 s* in ***N-paradigm***), and each trial was initiated by the animal refraining from licking the waterspout for at least 0.5 s. In the N-paradigm, trial initiation began with the water pump turning on, leading to licking before the presentation of the task sound. However, in the P-paradigm, a large drop of water was delivered *only* at the beginning of a task session, animals consumed the *free* water and then refraining from licking to initiate the first trial.

The daily training session ended naturally when the animals were no longer thirsty and did not lick the waterspout for three consecutive trials with presence of the water stream in the ***N-paradigm***, or for three target trials in the **P-paradigm**.

### Training Procedures and Stimulus (Training) Sets

Animals were trained on two tasks - to either categorize pure tones or amplitude modulated stimuli. The *stimulus sets* used for training the two tasks included two general sets of stimuli: one set consisted of pure tone stimuli that varied along the frequency dimension (***TN-task***), and the other set consisted of amplitude modulated noise stimuli that varied along the temporal dimension (***AM-task***). In the ***TN-task***, the initial stimulus set in the ***N-paradigm*** consisted of 6 stimuli (***Table-1***) that spanned a ∼7 octave frequency range with ∼1.5 octave increments between the pure tone stimuli. The initial training sets were slightly different in the **TN-task** for the animals trained with the ***P-paradigm***. For those animals, the initial **TN-task** training set consisted of 15 pure tone stimuli with varied frequency range. In one ferret (***Ganges***) the stimulus frequency range was also ∼7 octaves as in ***N-paradigm*** training. The other two ferrets trained with the ***P-paradigm*** (***Nile and Bramble***) were given an initial training set with finer frequency resolution that spanned only a ∼2.5 octave frequency range with ∼1/6 octave increments between adjacent stimuli (*see **Table-1***). The initial training set in the ***AM-task*** for both paradigms consisted of 6 stimuli ranging from 4-60 Hz amplitude modulation (*100% depth*) using white noise as carrier (***Table-1***). Nevertheless, the stimuli from every set were divided into three regions along the relevant feature dimensions (*Spectral and Temporal*), in which stimuli in the *Middle* region were labeled as ‘***Target’*** stimuli and stimuli in the two flanking regions (*Low and High*) were labeled as ‘***Reference*** (non-Target)’ stimuli (***Table-1***). As indicated above, in the ***P-paradigm***, animals learned to lick the waterspout for water reward if the stimulus was a member of the “***Target***” sound category, and to refrain from licking to stimuli in the “***Reference***” sound category. In contrast, in the ***N-paradigm***, to avoid a mild electric foot/tail shock, animals learned to refrain from licking the waterspout if the stimulus was a member of the “***Target***” sound category, but could lick freely for ***Reference*** stimuli (see ***Figure 1***).

*The training procedures* were similar to our previous work and have been fully described in prior publications (***Yin et al 2016, 2020***). In brief, task training began after a 1-2 day habituation period, during which the animals became familiar with the testing environments (waterspout, speaker, light configuration, testing box dimensions etc.) and were given positive reinforcement (liquid reward) in the behavioral training arena. For both behavioral paradigms, all ferrets were first trained on the three-range categorization task along the frequency feature dimension (***TN-task***) with the frequencies listed in ***Table-1***. After reaching behavioral criterion, three of them (see **Table-1** for details) were further trained on the task along the temporal dimension by using a set of 6 amplitude-modulated noise stimuli, and learned to perform the three-range categorization task now based on modulation rate (***AM-task***). Intensity cues (initially ∼40 dB louder for Target over Reference sounds) were used to direct the animal to attend the contrast between the sound categories at the earliest stage of TN-task training. These intensity cues were gradually reduced and eventually completely removed over the course of training. No intensity cues were necessary or used for **AM-task** training as animals had already learned the basic task structure following prior ***TN-task*** training. All 5 ferrets successfully learned to categorize task stimuli to behavioral criterion, based on the trained feature dimensions (frequency in the ***TN-task***, and AM rate in the ***AM-task***), and were able to generalize their performance to novel stimuli along the trained dimension (some of these behavioral data have been previously published - in ***Yin et al 2016***).

### Behavioral Assessment and Psychometric Functions fitting

#### Behavioral assessment

Behavioral performance of animals performing the Go-NoGo tasks was assessed based on analysis of their stored digitized lick signals (the TTL signal) to task stimuli during daily task sessions. The timing of the rising edge of the first-lick (***FL***) after each sound presentation in the ***P-paradigm***, or the falling edge of the last-lick (***LL***) *before the end of the shock window* in the **N-paradigm,** were recorded as the behavioral *response* in a trial (*illustrated in* ***Fig. 1A-B***), or in the absence of a lick, as *no-response*. A *response* that fell *within* a time window (0.25-2.0 s after sound onset) in the ***P-paradigm*** or fell *before* the shock window (0.85-1.25 s from sound onset) in the ***N-paradigm*** was defined as a *Hit* if the sound was a ***Target*** sound, or a *False Alarm* if the sound was a ***Reference*** sound. Otherwise, if the *response* occurred *after* the time window in the ***P-paradigm*** or occurred *within* the shock window in the **N-paradigm**, it was defined as a *miss* after a ***Target*** sound or a *correct reject* after a ***Reference*** sound. The hit rate (***HR***), false alarm rate (***FAR***), miss rate (***MR***) and correct reject rate (***CR***) were computed in each behavioral session. Two metrics used for assessing animal behavioral performance were derived in relation to ***HR*** or ***FAR***. One was the discrimination rate (***DR***), was computed as **DR = HR*(1-FAR)*100%** (***Fritz et al 2003***). The other measure used was the ***d’*** measure as *d’ = Φ^−1^ (HR) – Φ^−1^ (FR)*, and Φ^−1^ denotes the *probit* transformation. Significant task performance on a behavioral session was defined as either or both ***DR***>=40% or ***d’***>=1. The criterion for task mastery was the achievement of significant performance in three consecutive training sessions, with more than 100 trials in each session.

*Psychometric Functions and Fitting*: the psychometric function (***PF***) describes the probability of the animal making a *response* as a function of the quantitative characteristic along the task-relevant stimulus feature dimensions (the frequency in the ***TN-task*** or the amplitude modulation rate in the ***AM-task***). We used a generic 4-parametters Logistic (sigmoid) ascending function to fit the lower boundary of the PF between low to middle range (***function 1A***), and the descending function for the upper boundary from middle to high range (***function 1B***):

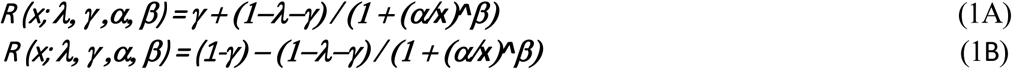

In these two formulas, the parameter ***γ*** corresponds to the *baseline rate* or *guess rate*, reflecting the situation where the animal is making a correct response purely by guessing, irrespective of the stimuli. The parameter ***λ*** corresponds to the *lapse rate*, reflecting the situation when the animal fails to respond correctly (making a *non-response*) which accounts for random errors regardless of the stimulus. The parameter ***α*** determines the reflection point of the PF function and reflects the border that divided the adjacent sensory regions. The parameter ***β*** is a measure of the sensitivity, and determines the slope (***k***) of the curve at the reflection point.

In the **TN-task, the stimulus frequency** was converted into log-scale in PF fitting. The fitting of the PF was derived using the MATLAB function ‘***fit’***. The fitting conditions were set by the MATLAB script ‘***fitoptions’***, in which the search for the optimal fit started with initial parameter conditions at [***γ****=0.5,* ***λ****=0.5,* ***α0****=middle point,* ***β****=1*], and the search ranges were confined between [0 1] for ***γ*** and ***λ***, between [*min max*] of stimuli quantity for ***α***, [0 100] for ***β***. The inflection point (*α*) of the fitted curve indicates the perceptual boundary between the *Reference* and *Target* sounds categories, and the difference between ***α*** and the middle point (***α0****, the boundaries*) can be used to quantity the possible bias (or shift, ***Δα****=****α****-****α****0*) of the boundary. The sensitivity of the ***PF*** at the boundaries could be described by the slope (***k***) at the inflection point derived from the parameter ***β***. The goodness of the fit is quantified by the cost function of normalized root mean square error (***NRMSE***), in which a perfect sigmoid fitting has a value of 1, and a zero or negative value indicates that the fit is no better than a straight line. The PF fitting was performed for each of the behavioral sessions. Following initial behavioral training and reaching behavioral criterion on all tasks, most of all subsequent behavioral sessions (*388/469=83%*) conducted during neurophysiological recording showed a good sigmoid fit, indicating successful categorical performance.

### Implant Surgeries and Behavioral Methods during Neurophysiological Recordings

After animals reached behavioral criteria for all trained tasks and attained stable performance, implant surgeries were conducted under aseptic conditions while the animals were deeply anesthetized with 1-2% isoflurane. A stainless steel headpost was attached to the ferret skull with bone cement (***Palacos MV +Gentamicin, Heraeus***). About 2-3 weeks later, following recovery from surgery, animals were gradually habituated to head restraint in a customized stereotaxic frame and retrained on a head fixed version of the tasks. The stereotaxic frame was positioned on a stable platform in a double-walled soundproof chamber. Acoustic stimuli were presented from a free-field loudspeaker (***Manger, German***) located ∼1.2 m directly in front of the animal. During neurophysiological recordings, all sounds were amplified (***MA-3, Rane***) and presented at 60-75 dB SPL. Animals needed several weeks to re-train and regain successful behavioral performance on the head fixed version of the tasks.

After the head-restrained animals re-attained behavioral criterion performance levels as detailed above, they were prepared for neurophysiological recordings. A small craniotomy (1-2 mm in diameter) were made over the target brain regions, and neurophysiological recordings commenced. Daily recordings were conducted with a 4-tungsten microelectrode array (*1-5 MΩ, **FHC***) simultaneously inserted into the brain through the craniotomy using an Electrode Positioning Drive (***EPS-drive, Alpha-Omega***). Raw neural activity traces were digitally acquired/stored by a commercial data acquisition system (***Alpha Lab, Alpha-Omega***). In each recording session, multiple single-units were isolated offline by customized spike-sorting software, based on a PCA and template-matching algorithm (***Meska-PCA, NSL***). Recordings were made over a period of 6–18 months in each ferret from two brain regions: dorsolateral frontal cortex (***FC***) and auditory cortex. The craniotomies were expanded as needed over the course of the recordings. Only FC data are included in this manuscript.

A typical behavioral neurophysiological session included at least one sequence of three recording epochs in which the animals were: 1) initially passively listening to a selected stimulus set (from the TN-task or AM-task); 2) then actively engaged in performing the selected task while being presented with the identical stimuli from Epoch 1; 3) passively listening once more to the same stimulus set. In each epoch, trials were generated in a pseudorandom order to make sure each of the listed stimuli were equally presented and repeated at least 10 times. Typically, the first behavioral session at each recording site/depth was the TN-task (with all three epochs). Following the first behavioral session, if the animal was still motivated to continue behavior and if the units were stable, then recordings continued with a second session with the animals performing the AM-task. Occasionally, the task order was reversed during recordings. Overall, however, there were more TN-task behavioral physiology sessions (***n=342***) than AM-task (***n=127***) sessions. In some recording sites, both TN-and AM-task (***n=119***) were available (*see **Table-2** below*).

### Neuroanatomical Methods for Localization of Recording Sites

The locations of neurophysiological recordings in the dorsolateral FC were situated ∼2-8 mm lateral to the midline, and ∼25-30 mm anterior to the occipital crest. This location corresponds to a frontal cortex area including both the anterior sigmoid gyrus/premotor cortex (ASG/PMC) and a more anterior dorsolateral FC region of the proreal gyrus (PRG), based on the ferret brain atlas (***Radtke-Schuller, 2018***). The craniotomy was gradually expanded over the course of 6-18 months of recordings, and in some animals a second separate craniotomy was made over the frontal cortex in the opposite hemisphere for additional recordings. Locations of the recording electrodes relative to two landmarks on the surrounding bone cement were noted for later alignment of all electrode penetration sites on the final craniotomy images and a map of all recording sites in each animal (*Supplementary Figure 3A*).

Following the completion of neurophysiological recordings in each animal, tracer deposits were placed in the FC recording sites to enable future histological confirmation of the recording areas. In each animal, two penetration sites in each identified cortical area were selected. In three of the ferrets (***Gong, Ganges and Nile***) iron electrolytic deposits were made in these sites. In the other two ferrets (***Guava and Bramble***) labeled marking was introduced by injection of a small volume (*0.3-0.7 μl*) of fluorescent dye (fluoro-emerald (***FE****) or fluoro-ruby (**FR**), 0.1 mg/ μl in saline) (**Molecular Probes Inc., Eugene, OR, USA**)*. The chosen recording sites for labeling were identified first by responses evoked during task performance. The iron deposits were made by passing a small constant current through stainless-steel electrodes (5-7.5 μA for 300 s). The ***FE/FR*** injections were made using a 5.0 *μl* Hamilton syringe with 30-45 degree tip. Following perfusion of the animals with a 4% paraformaldehyde solution, and subsequent histology, confirmation of injection sites in 50 *μm* coronal brain slices of the fixed ferret brains was achieved using a Prussian blue reaction to highlight the iron deposits, or by identifying the fluorescent deposits by directly inspecting with a fluorescent microscope. Recording sites were successfully identified, confirming localization in dorsolateral ***FC*** (including ***dPFC*** and rostral part of ***ASG***/***PMC****, see* in ***Supplementary* Figure 3A**) based on the ferret brain atlas (***Radtke-Schuller, 2018***).

### Methods for Neuronal Data Analysis

The single units isolated from FC recordings were first evaluated by examining responses in three time epochs relative to stimulus onset: a) pre-onset baseline (-*100 ms pre-onset – i.e. 100 ms before stimulus onset*), b) onset (*stimulus onset to 250 ms post-stimulus onset*), and c) sustained (*250 ms post-stimulus onset to* the full stimulus duration). Neuronal activity in each of these three time epochs was analyzed with respect to sound category (***Reference*** or ***Target***). Neurons that showed significant modulation in at least two of the following six testing pairs were included in the final data analysis: 1) onset response vs baseline response (***Reference***), 2) onset response vs baseline response (***Target***), 3) ***Reference*** vs ***Target*** response (onset), 4) sustained vs baseline response (***Reference***), 5) sustained vs baseline response (***Target***), 6) ***Reference*** vs ***Target*** response (sustained). A total of *1480* neurons isolated from ***350*** recording sessions over a period of ∼12-18 months passed these screening criteria and were therefore included for further analysis (see ***Table-2*** for details). The basic neuronal metrics applied included *categorical index* (***CI***), discrimination index (***DI***), *Choice probability (**CP**)* etc., and were described in previous studies (***Yin et al 2020***).

#### Clustering Neurons based on the Mean Response Profiles

Neuronal activity in FC regions often conveys information about attention, sensory input, stimulus category, decision making, and motor planning. We sought to analyze these correlations of neuronal activity with internal states and behavior by using principal component analysis (***PCA***) to extract the major components conveying the dynamics of the response profiles evoked by stimuli. The neurons were then partitioned based on the weights from the dominant components using K-means clustering.

The details of this analysis are shown in a Supplemental figure (***Figure S5***). At first, the response profiles (***PSTHs***) to reference or target sounds category were averaged from all correct trials, and then smoothed with a Gaussian kernel (*100 ms* in length with *20 ms* standard deviation) with a *5 ms* moving step. The obtained profiles were normalized by subtracting baseline activity (obtained by averaging over 100 ms pre-stimulus period) and then scaled between [-1 1] by dividing the absolute maximum of the subtractions. Neurons with more than 10 correct trials in both reference and target trials were selected to form a 2D data matrix (time bins x neurons). In order to extract the dynamics for overall response profile, the portion of the data matrix from −100 ms pre-stimulus onset to 1000 ms post-stimulus onset was selected to perform PCA decomposition (without centering the columns of data since it had been standardized) in which the rows of the data matrix correspond to observations (the time) and columns correspond to variables (*neurons*) (***Supplementary Figure S5A***). The obtained PCs reflect the composition of the dynamic profiles in descending order in terms of the percentage of the total variance explained by the component (***Supplementary Figure S5B***). The profile contributions in the neuronal response could be estimated by the Least-Squares Solution of a linear equation:

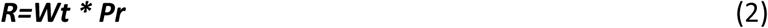

Where ***R*** denotes the response matrix evoked by Target or Reference (as in ***Figure S1A***); the ***Wt*** denotes the profile weights for each neuron; ***Pr*** denotes the extracted PC profiles which are scaled between [-1 1].

The majority of explained variance (65-84% of total variance across task variants) was captured in the first three PCs, which also showed remarkably similar profile pattern across task variants. We then partitioned the neurons into neuron clusters by using the K-means clustering method (Matlab function ‘***kmeans’*** with ***Squared Euclidean distance,*** *5 time repeats*) based on obtained ***Wt****’s* to profiles 1-3 (***Supplementary Figure S5C***). Three major clusters were confirmed, one cluster with transient responses and two clusters with sustained average response profiles (***Supplementary Figure S5D***). This analysis procedure is applied to the data matrix for different stimulus types (the ***Reference*** or ***Target****, or the combined*), for different behavioral contexts (passive or *active*), and for the neurons in each task (***TN*** or ***AM***) and Paradigm (**P**, **N**). Since FC neurons were most strongly driven by active engagement in the task and by Target sound category, we elected to use only the dataset from Target stimuli to extract the PC profiles and partition the neurons into clusters. The combined dataset (including responses to both Reference and Target stimuli) returned comparable PC profiles in which the explained variance by the first 3 PC profile was slightly decreased (by ∼1.8% in average across tasks).

#### Decoding Reference and Target Based on the Population Neuronal Activation Patterns

Averaged response of the neuron population (i.e. the PSTHs) could describe significant information about the stimuli (***Bagur et al 2018; Foffani & Moxon, 2004***). However, the activities induced by Reference and Target sounds in same FC population reflected distinct signals that played different functional roles during task performance, such as categorizing the stimuli, decision making/action selection and execution/maintenance of selected actions. According to the temporal stability hypothesis (*Jagadisan & Gandhi 2022*), these multiplexed signals might be aligned with distinct temporal structures of population activity patterns. In this way, the population temporal structure could supplement the rate code (the averaged response) in disentangling the different processes related to stimuli, decisions and actions over the course of time during behavior. Hence, we adapted the method to decode task stimuli based on the population temporal structures.

To simplify the analysis, neurons with more than 12 correct trials for each sound category (both ***Reference*** and ***Target***) were included in this analysis. We organized *first* a 3D pseudo-population data matrix (population trials x neurons x time bins), in which each population trial was constructed by randomly picking a correct ***Reference*** or ***Target*** trial from each neuron. To remove a potential overtraining effect, the data matrix was limited to include only 12 Reference and 12 Target population trials to make sure that no trial was re-chosen in each population draw. To assess temporal structure, the resulting pseudo-population data matrix was *then* normalized by its Euclidean norm (equivalent to its magnitude) to form a unitary vector in multi-dimensional neuronal space at each time instant during a population trial (*Jagadisan & Gandhi, 2022*). Thus, the normalized data matrix reflects a unity length population vector that points in an n-dimensional direction at a given time instant in each population trial.

Assuming that the population trials from Reference and Target were aligned with distinct temporal structures associated with different task events, a leave-one-out cross-validation (**LOOCV**) procedure was used to decode the trial type (Reference or Target trial) based on the stability of the temporal structure (as indicated in the flow chart in **Figure 5A)**. During the training stage, the template vectors for Reference (***V****ref*) and Target (***V****tar*) were generated by averaging 11 Reference or Target trials at a given time instant, with one Reference/Target trial left out as testing trial. The temporal stability (***S(****t, t0)*) of the population activation at the time instant (*t0*) was then quantified by the dot product of the template vectors (***V****ref* or ***V****tar*) and the single trial test vector (***V****test*(*t, ref/tar*)) at each time moment of the trial:

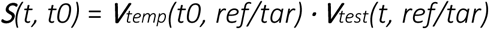

A testing trial with a larger stability metric (***S****(t, t0)*) produced from the Target vector (***V****tar*) was then classified as a Target trial or Reference trial. This procedure was repeated 12 times to make each trial serve as a testing trial for a given population data matrix draw. There was a total of 100 population draws in each of the task variants, which yielded a total of 2400 testing trials (1200 Reference and 1200 Target population trials). The temporal stability metric of the testing trials can be visualized in a 2D space constructed by Reference and Target vectors. As shown in ***Figure 5B*** for a few example snapshots at different time instants, test population trials were segregated to align to Reference and Target temporal structure over the course of time.

#### Extracting β-band rhythms from Local Field Potential (LFP)

Local field potential (LFP) data were collected during the same physiological recording session as single-unit data. The raw LFP data were first passed a notch filter to remove 60 Hz noise, followed by a high-pass filter at 1 Hz (2^nd^ order Butterworth) to remove slow drifts, and then down-sampled to 500 Hz. For those trials received electric shock in N-paradigm animal, the data around the shock period (∼400 ms) were set to zero. All trial traces were corrected by the averaged pre-stimulus level (100 ms before stimulus onset) to further remove the slow drifts on a trial-by-trial basis.

To extract the time-frequency (**TF**) power, the preprocessed LFP data were convoluted by a family of complex Morlet’s wavelets (***w(t, f0)***), which have a Gaussian shape both in the temporal (***α****t*) and frequency (***α****f*) domain around its central frequency (***f0***):

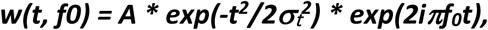

Where ***α****f **= 1/(2******ν α****t)* and a normalization factor ***A = (α****t* ***ν)^-1/2^*** make total energy equal to 1. A wavelet family is characterized by a constant ratio (***f0/α****f*). In our analysis, the wavelets we used were defined by *f0/****α****f* = 5. The analyzed frequencies ranged from 10 to 115 Hz with 50 equal log-spaced bins.

The **TF** power for each LFP trial were computed by the squared amplitude of the resulting complex spectrum. The power values were then converted to percent occupancy of the total power for each analyzed frequency channel at each recording site. The TF power data were then averaged across trials for different trials types, such as Reference *vs* Target in terms of stimulus type, or Hit/Miss *vs* Correct reject/False alarm in terms of task behavior), and standardized by working along time and frequency dimension. The resulting TF powers pooled across all recording sites for statistical analyses.

---

**Supplementary Figure 1.**
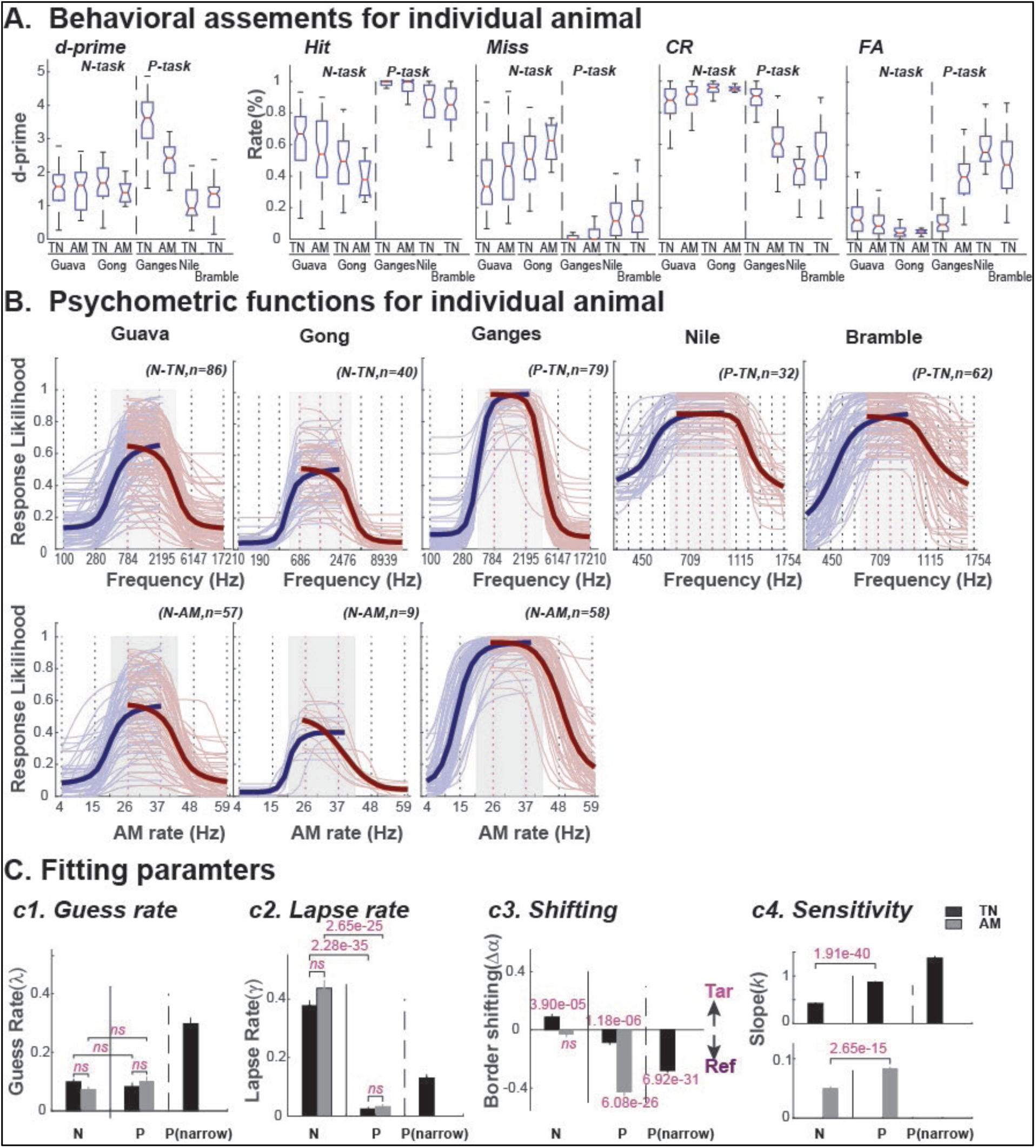
Behavioral Assessment Metrics and Psychometric Functions (PF) for Individual Animals. A. **Boxplots illustrating the performance assessment metrics** for each individual animal and task. The analysis included all behavioral sessions during physiological recordings. Going from left to right, the plots show the ***d’*** (behavioral performance) and the reaction rates for each trial types: ***Hit***, ***Miss***, *Correct Reject* (***CR***), and *False Alarm* (***FA***), respectively. In general, animals trained on the N-paradigm (***Guava*** and ***Gong***) made more behavioral errors as a result of the *Miss* trials (by failing to *stop licking* after a ***Target*** sound) than as a result of *False Alarm* trials (by mistakenly stopping *licking*, after a ***Reference*** sound), in both the **TN** and **AM** tasks. However, by contrast, animals trained with the P-paradigm (***Ganges***, ***Nile*** and ***Bramble***) often made more False Alarms errors (mistakenly licking *the spout* after a ***Reference*** sound) than Miss errors (failing to *lick the spout* after a Target sound). This opposite pattern of error types between the ***N-*** and ***P-paradigms*** likely reflects different strategies developed during training to maximize water intake. In P-paradigm, this would entice the animals to maximize their water intake at the cost of experiencing a 3-5s timeout for licking after a Reference stimulus (leading to more FA errors). In the N-paradigm, this would entice the animals to lick after Target stimuli and hence tolerate a mild shock (leading to more Miss errors). This tendency was accentuated when tasks became more difficult, such as in the **AM** task (ferrets: *Guava* and *Gong* in ***N-paradigm***, and *Ganges* in the ***P-paradigm***) or in the **TN** task with finer resolution (*Nile*, *Bramble* in the ***P-paradigm***). B. **The PFs obtained from each of the behavioral sessions** by fitting the likelihood of the behavioral responses after each of the stimulus vs stimulus parameters (frequency in the **TN** task or modulated rate in the **AM** task) with a 4-parameter logistic function (***see Methods section: Behavioral Assessment and Psychometric Function fitting***). Two fittings were made for each behavioral session: from Low to Middle (thin blue lines), and from Middle to High (thin pink lines). The thick blue and pink lines show the averaged PFs across all sessions for each animal/task. C. **Bar plots depicting the extracted parameters** from the fitted PFs grouped by training paradigms (**N** or **P**) and stimulus sets (**TN**, **TN**-(narrow) and **AM**). The data were averaged from the two fits of Low-to-Middle and Middle-to-High. In a 4-parameter logistic function, the Lapse Rate (***γ***) estimates the likelihood of the animals failing to make the learned behavioral response to a **Target** stimulus (to *stop licking the spout* in the ***N-paradigm*** or initiate *licking of the spout* in the ***P-paradigm***) even when the stimulus is far above the threshold of the PF (***Miss*** *errors),* which often reflects inattention or motor errors. The Guess Rate (***λ***) estimates the likelihood of an erroneous response to a **Reference** stimulus when the stimulus is far below the threshold of the PF (***FA*** *errors*), which likely reflects a random process of pure guessing. These estimated likelihoods are closely correlated to the lower or upper asymptotes of the fitted PFs. Thus, the combination of Lapse and Guess shift the PF up or down reflecting a different default (or habitual) behavioral strategy that was developed over the course of training (***c1-c2***). Low Guess with high Lapse rate causes a *downward-shift* of the PFs (as seen in all **N-paradigm** animals for both **TN** and **AM** tasks, and in **1B** for individual **PF**s), which reflects a *conservative* strategy or disengagement. A high Guess with low Lapse causes an *upward-shift* of the PFs (as seen in **P-paradigm** animals, but more striking in **P(narrow)** animals in TN-task, **Nile** and **Bramble**; also seen in **1B** for individual **PF**), which reflects a *liberal* strategy or improved vigilance. The behavioral response errors could also be associated with shifting of the PFs (**Δα**), as measured by the subtraction of the reflection point (**a**) of the PF from the midpoint (**α0**) between **Reference** and **Target** (*on a logarithmic scale*). Negative border shifts indicate the PFs shifted into **Reference** zone, causing more *FA errors* to Reference stimuli. Positive border shifts indicate the PFs shifting into Target zone, thus causing more *Miss* errors to Target stimuli (***c3***). Animals in the ***N-paradigm*** exhibited either a minor *positive shift* (in the **TN** task) or no shifts (in the **AM** task) of the PFs. Animals in the ***P-paradigm*** showed significant *negative shifts*, more robust in the **AM** task and TN-task with finer resolution (***P(narrow)*** groups animals: ***Nile*** and ***Bramble***). The different behavioral strategies also reflect the sensitive shift of the PFs between N and P-paradigm (***c4***). The numbers above the bars are the *p-values* from either *Two-sample t-test* (***c1-2,c4***) or *paired t-test*(***c3***), *ns* indicates no significance of the testing (p<0.05).

**Supplementary Figure 2.**
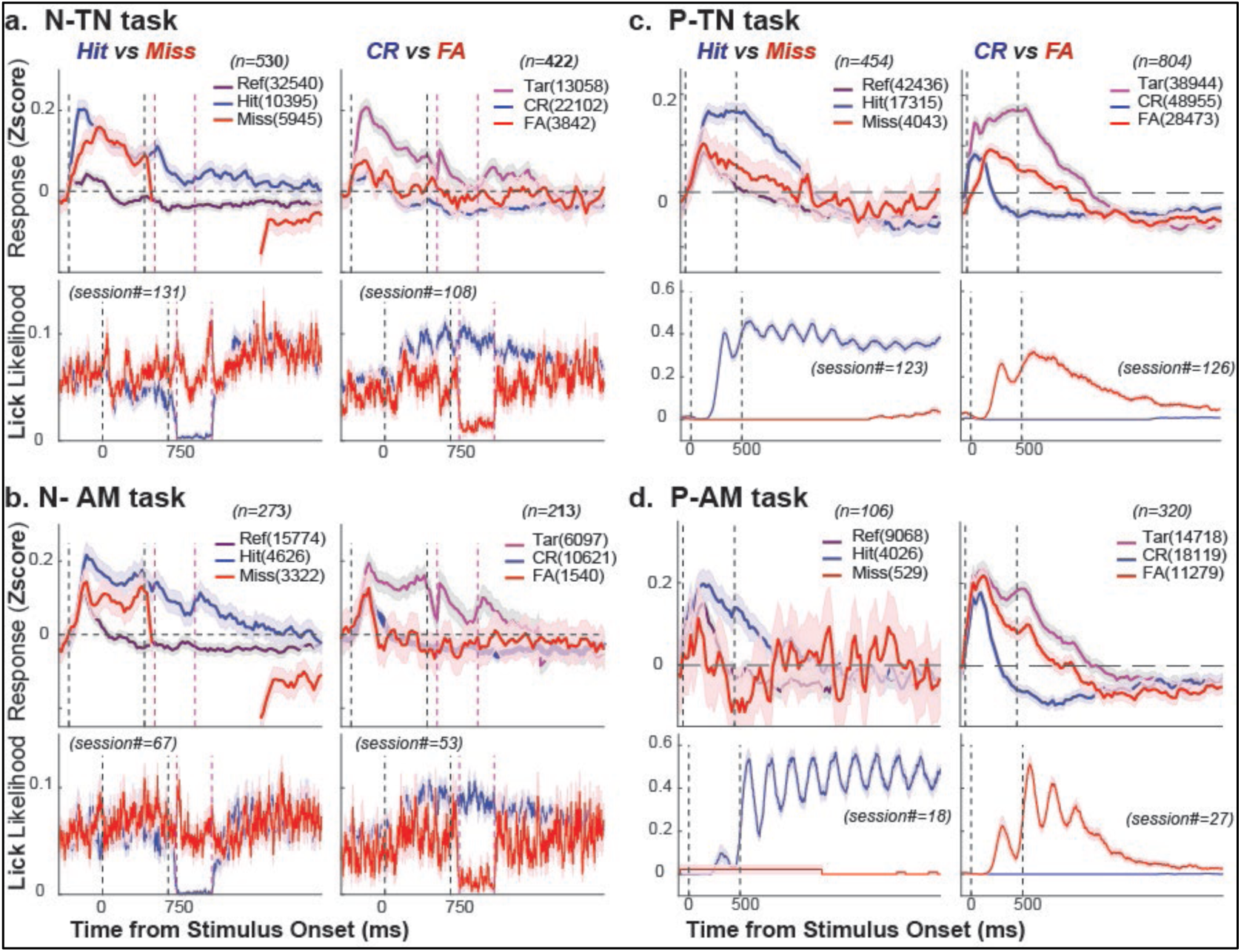
Neuronal Correlates of Behavioral Choice – Analysis of Error Trials. The neuronal correlates of behavioral choice were analyzed on a trial-by-trial analysis basis. In each neuron, the spiking activity on each behavioral trial was binned in 100 ms sliding windows with 5 ms steps and then standardized across all trials and time bins (a total 105 time bins beginning 100 before stimulus onset and up to 2500 after stimulus onset). The normalized activities of all neurons with at least 2 trials in selected trial types were pooled together to form a large 2D data matrix (*trials* x *time bins*) for the corresponding analysis of trial types. The top row of panels in each plot showed the grand mean and the 95% confidence interval for the mean (*indicated by the shaded area around line*) of activities from the selected trial types: ***Hit*** vs ***Miss*** (left panel), and *Correct reject* (***CR***) vs *False alarm* (***FA***) (right panel). The total number of trials pooled from the population are shown in the parentheses following trial type legend in each panel. The averaged licking profiles for different trial types were shown in lower row of panels in the plot. The black vertical dashed lines indicate stimulus onset and offset times, and the red vertical dashed lines in (***a, b***) mark the onset and offset of the shock window in N-paradigm. There is a marked interesting difference between the N- and P-Paradigm responses to the error trials. In the N-paradigm (***a***, ***b***), in response to **Target** stimuli, there was no significant difference between the averaged PSTH of ***Miss*** *versus **Hit*** trials (*panels on the left*). The neuronal responses to **Reference** stimuli were also similar whether evoked by **FA** or **CR** trials (see *panels on the right*). However, in the P-paradigm (***c***, ***d***), the incorrect trails (**Miss** and **FA**) showed different response profiles in comparison with the response evoked by correct trials (**Hit** and **CR**). This contrast between the N- and P-paradigms likely correlates with the different behavioral strategies in the two training paradigms. In the N-paradigm, as shown in the ***Supplementary* Figure 1C**, most ***Miss*** Errors were attributed to inattention or motor errors (as estimated by *Lapse rate.* See **Methods**), reflecting a *conservative* strategy or disengagement. As shown in the licking profiles in lower panels, ***Hit*** and ***Miss*** yielded a similar trend of licking profiles before the shock widow, it seemed that animals made a correct initial reaction after target stimuli but resume the licking earlier in time. Therefore, the similar response profiles between Hit and Miss (neuronal decision) indicate a mismatched behavioral outcome. In the P-paradigm, a significant portion of errors arose from the border shifting into Reference of the PFs (***Supplementary* Figure 1B*, 1C***), especially in the more challenging tasks such as the ***AM****-* or ***TN****-tasks* with finer rate or frequency resolutions, reflecting a *liberal* strategy or improved vigilance. The difference of the neuronal response profiles between the CR and FA (neuronal decision) were faithfully reflected behavioral outcome which is consistent with the choice probability measures (see Supplementary Figure 3).

**Supplementary Figure 3.**
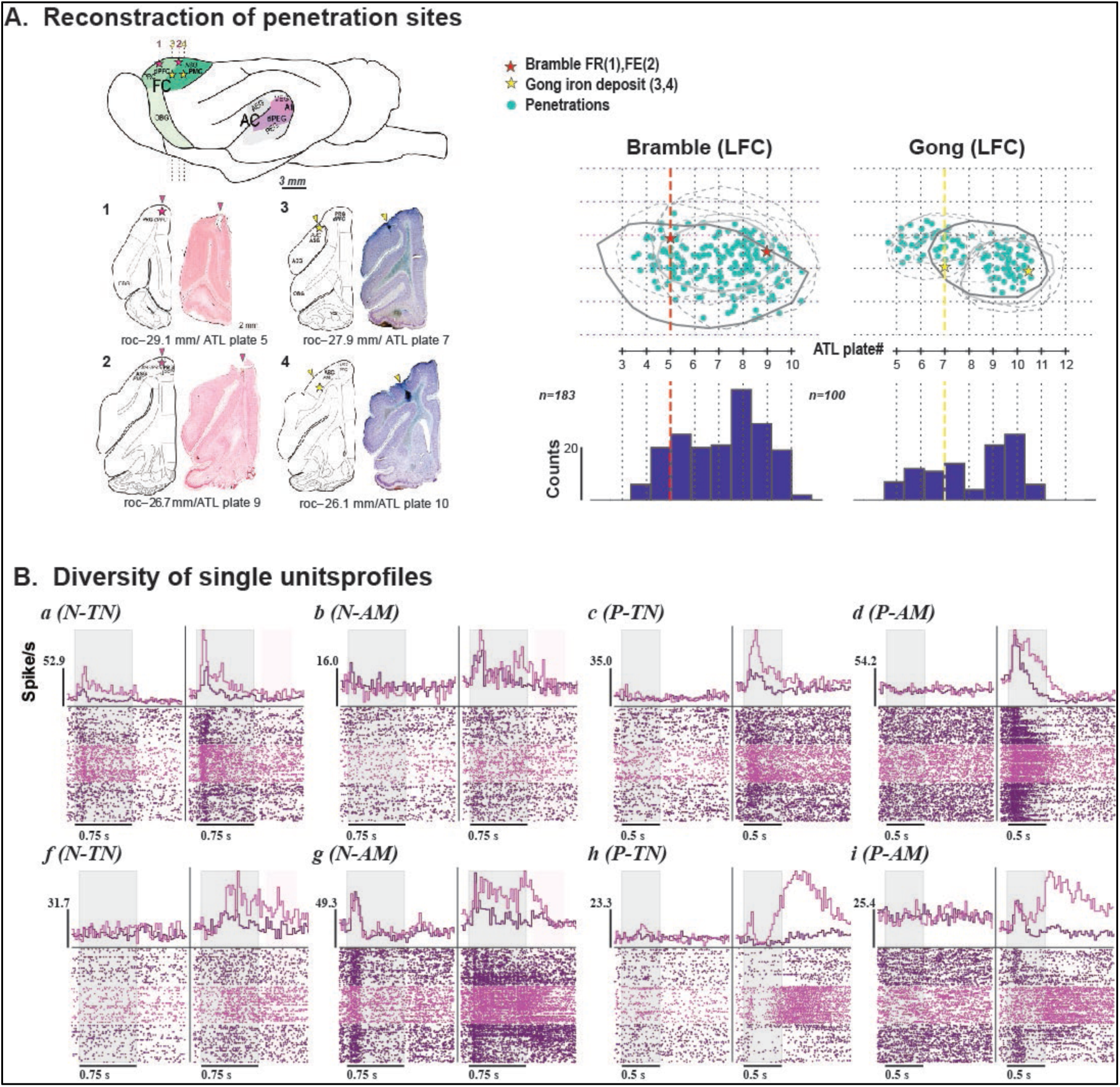
Localization Of Recording Sites and Diversity of Single-Unit Response Profiles**. A.** Electrode recording sites are highlighted on the surface of a sideview of standard ferret brain. Stars in frontal cortex (**FC**) indicate the fluorescent dye (*yellow*) injections sites in one of the N-paradigm animals (***Gong***), or the iron deposits (*red*) from one of the P-paradigm animals (***Bramble***). The coronal sections across each of the marked sites (***1-4***) are displayed below with the corresponding atlas section (***Radtke-Schuller, 2018***) to determine the locations within these areas. The four marked sites were histologically confirmed to be span from the **dPFC** to rostral part of **ASG**/**PMC**. The scatter plots on the right are shown the electrode penetrations and the marked sites overlaid on photo view of the initial and final craniotomy (*the solid lines*) and its expansions (*the dashed lines*). The barplots below each of the scatterplots are aligned to the recording penetrations along the rostral-caudal atlas plate, which confirmed that the majority of our recording sites were in ***dPFC*** and rostral ***ASG/PMC***, between plates 5-10. **B.** Responses of FC neurons were modulated by the experimental contexts (*passive* vs *active*) and stimulus types (***Reference*** or ***Target***). Most neurons displayed a more robust response in the active state, and a weaker or no response in the passive state (panel ***b-f, h-i***). However, some neurons did show moderate responses to stimuli in the passive state (***a, g***). **Target** stimuli usually evoked a sustained response as in panels (***f, g***), or mixed profiles comprised of an early transient and late sustained excitatory (panels ***a-c***, ***h-i***) or inhibitory (panel ***d***) response patterns. **Reference** stimuli usually evoked moderate transient response (panels ***a-d, g, i***) or weak/no response (panels ***f, h***).

**Supplementary Figure 4.**
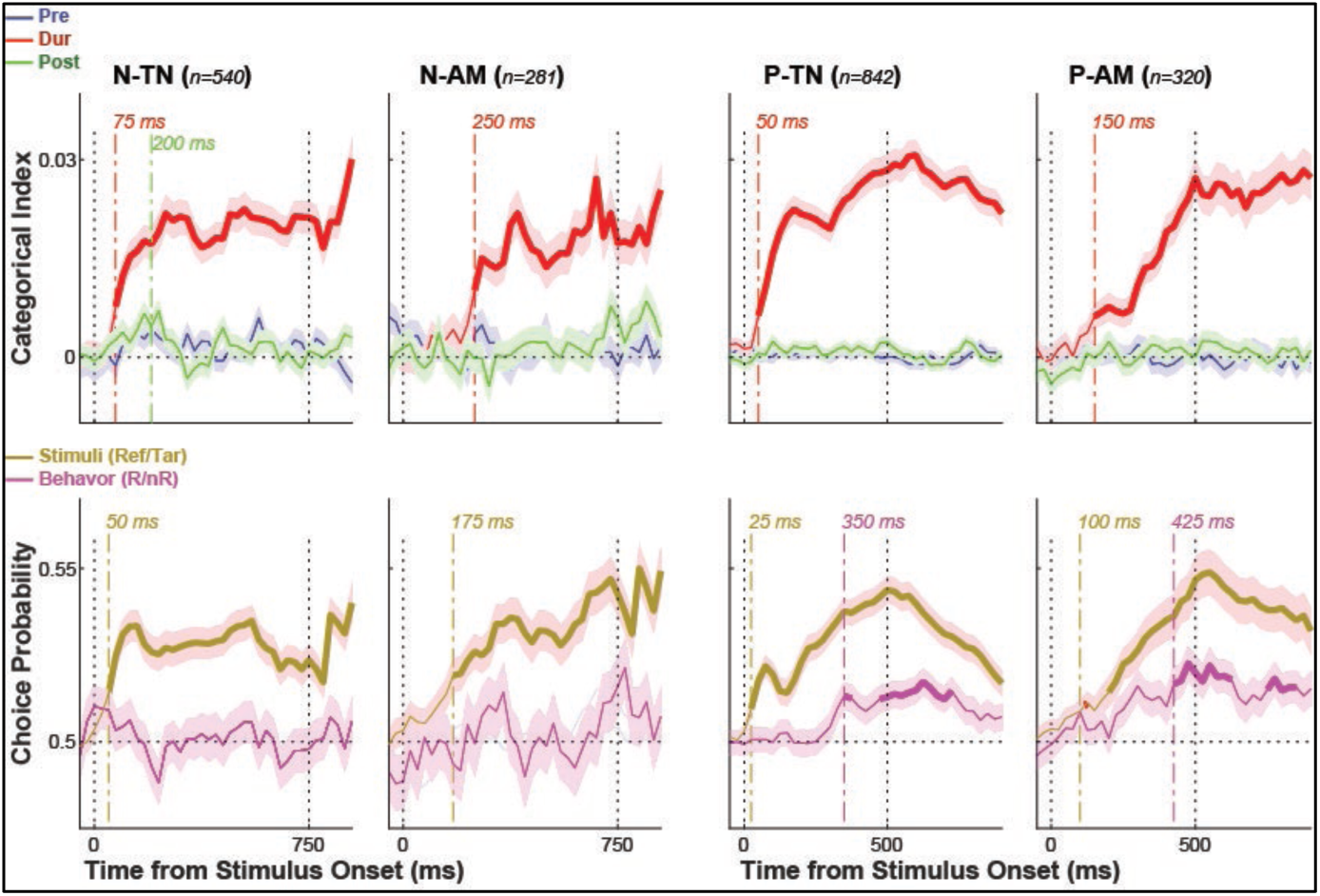
The dynamics of categorical information (CI) and the choice probability (CP) in FC neurons across the task variants (from left to right: N-TN, N-AM, P-TN and P-AM respectively). *<u>Top row</u> of panels* show the dynamics of the categorical information (extracted from 100 ms sliding time window with *25* ms step) about the stimuli (***Reference***/***Target***) during animals actively engaging in the task (*red*), and passively listening to task stimuli pre/post-task (*blue/green*). Not the similarity of the dynamics across different training paradigms/task. The significant categorical information about the stimuli could appear as early as 50∼75 ms after stimulus onset in TN-task, it appears much late in AM task (150∼250 ms). *<u>Bottom row</u>* of panels shows the likelihoods of a neuron preferring a Target over a Reference stimulus, or a Response (Hits for Target + False alarm for Reference) over a non-Response (Miss for Target + Correct reject for Reference) behavioral choice during engagement in a task performance. Note that, in all task variants, FC neurons are driven by the stimulus far more than by behavioral choice. The information related to behavioral choices are relatively weak. For instance, only trace amounts of information about the behavioral choice appear in the P-paradigm (both TN- and AM-task), and there is no significant information observed in the N-paradigm (both TN- and AM-task). <u>In all panels</u>, the vertical dotted lines indicate stimulus onset and offset times, and the vertical dash-dot lines mark the times when CI/CP show significant non-zero value (p<=0.05 by One-Sample t-test with Bonferroni correction of repeat measures). The bold portions of the curves indicate the portions with significant non-zero values.

**Supplementary Figure 5.**
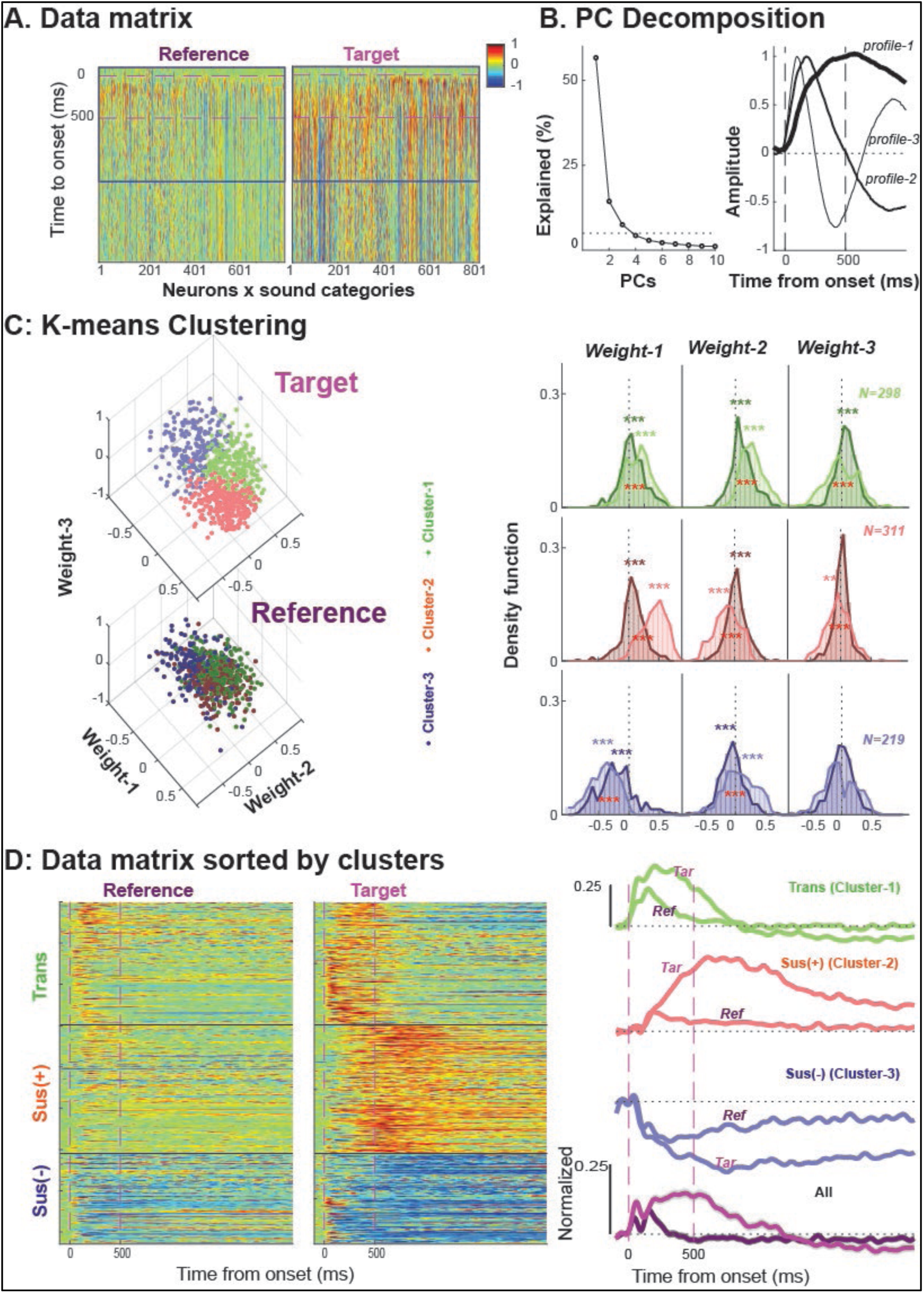
Methods for PCA and clustering. (**A**) **Data matrix:** An example Data matrix of the combined response profiles to **Reference** (left) and **Target** (right) stimuli across all neurons (including all correct trials) collected during task engagement. The response profiles were corrected by baseline firing rate (averaged over 100 ms pre-stimulus period) and scaled between [-1,1] by the maximum of the absolute value). The horizontal magenta dashed line indicates stimulus onset and offset times. PCA was applied to the population data matrix for Target (or in combination with Reference) sound category, with the time (from −100 to 1250 ms, 5 ms steps) as observations (highlighted by the blue rectangle), and the neurons as the variables. **(B**) **PC decomposition.** The line plot on the left shows the percentage of explained variance of the first 10 principal components. The first three PCs explained over 75% of the total variance of the response dynamics. The plots on the right are the three response profiles derived from the corresponding PC1,2,3 by normalizing by their absolute maxima. (**C**) **K-means clustering.** The profile contributions or weights (***wt***) in the neuronal response could be estimated by a Least-Squares Solution of a linear equation, **R = wt * Pr**, where **R** denotes the averaged response evoked by Target or Reference for each neuron, **Pr** denotes the extracted profiles. The neurons were partitioned into three major clusters by K-means clustering based on the obtained weights corresponding to profiles 1,2,3. The scatterplot on the left illustrates the representation of the neurons in the derived space to **Target** (top) and **Reference** (bottom) sounds. The plots on the right show the distribution of the *weights* from Reference (dark color) and target (light color) in each cluster. The stars above each distribution indicate weights with significant non-zero means (by one-sample t-test). The stars between the distributions indicate a significant difference between the weights from Reference and Target (by *paired t-test*). *, **, *** correspond to *p-*values< 0.05, 0.01, and 0.001 respectively. (**D**) **Data sorted by clusters**. The different clusters show distinct response profiles. The heatmaps on left are the same Data Matrix in (**A**) when rearranged by the neuron clusters. ***Cluster-1*** neurons (*green*) often have early responses after stimulus onset, quickly reaching a peak, and lasting a relatively short time, and hence is referred to as a transient (***Trans***) response. ***Cluster-2*** neurons often lack onset responses, showing instead a late buildup and long-lasting sustained excitatory response profiles (***Sus(+)***). ***Cluster-3*** neurons display a long-lasting suppression (***Sus(-)***) of activity that begins after stimulus onset. These distinct response profiles are evident in responses evoked by both **Reference** and **Target** stimuli but are more robust in **Target** responses. The line plots on the right depict the averaged response profile to **Reference** and **Target** stimuli grouped in terms of the three (green, red, and blue) clusters. The grand averages of the overall population are also shown in the bottom panel.

**Supplementary Figure 6.**
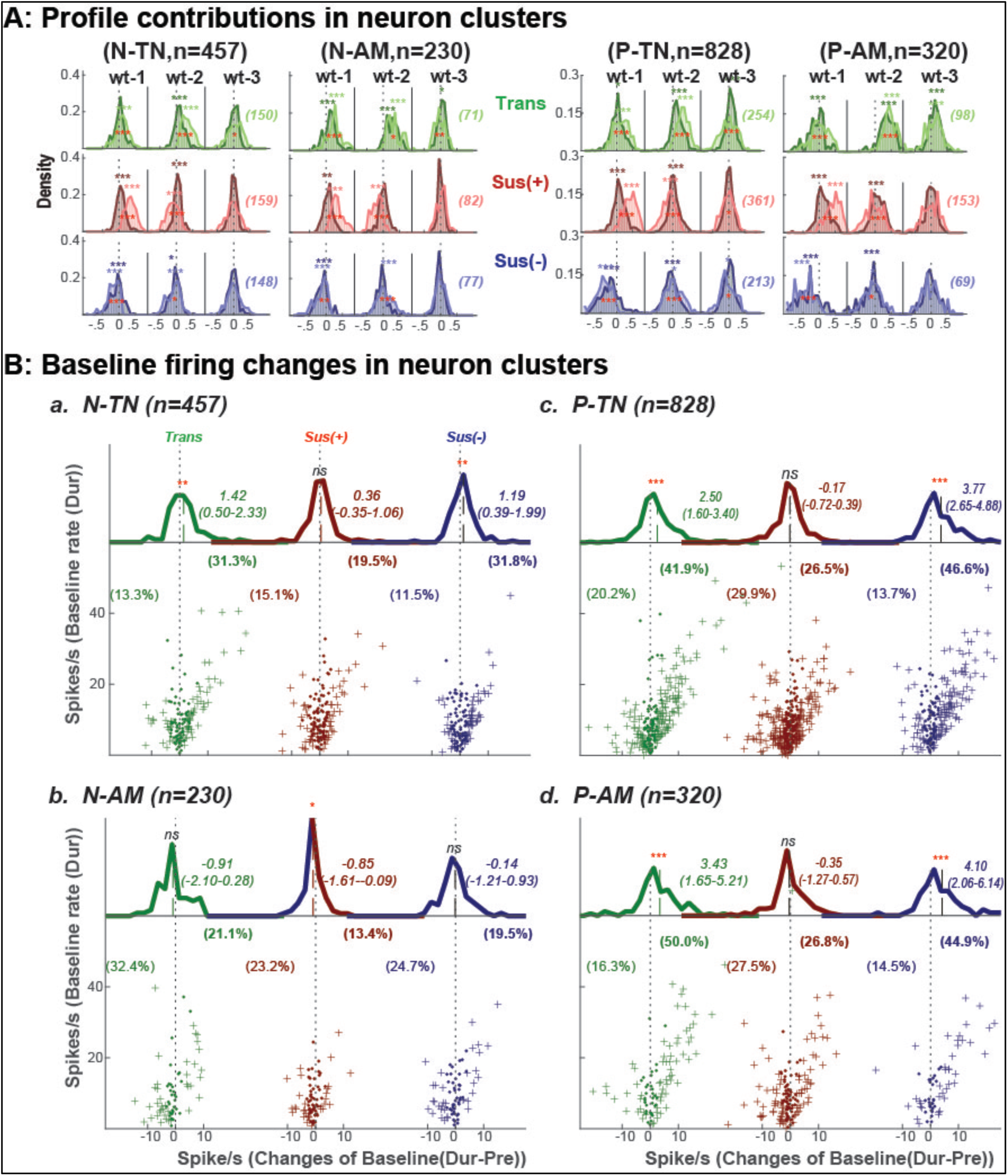
Profile contributions and baseline firing in FC clusters. **(A) Three similar temporal profiles** were extracted from the population response dynamics across different task variants as depicted earlier in **Figure 4B**. Their combination yielded three neuron clusters. The bar plots show the density distribution of the weights corresponding to the profiles in the each of FC neuron clusters in the different tasks. In all task variants, the ***Trans*** cluster weighted more from profiles 2,3, while the two sustained clusters weighted mainly from profile-1 (with either positive (***Sus(+)***) or negative (***Sus(-)***)) contributions. The averaged weights for **Target** (*lighter color*) were usually greater than in Reference (*the darker color*). **(B) Activity of FC neurons** was strongly modulated by task context. As shown in ***Figs 2 & 3***, most FC neurons displayed little or no response while passively listening to task stimuli but were strongly activated when animals actively engaged in a task. The modulation of neuron activity by task context (***passive*** vs ***active***) also occurred in the baseline firing rate of some **FC** neurons (neuron example ***e*** - ***f*** in ***Figure 2***). The four panels here show modulations of baseline firing rate in relation to the neuron clusters obtained from PC clustering analysis (see ***Figure 4***). The *bottom* scatter plot in each panel illustrates the relation between baseline firing-rate changes (*x-axis*) and the baseline firing-rate in the active state (*y-axis*) in each of the three neuron clusters (***Trans*** - *green*, ***Sus(+)*** - *red* and ***Sus(-)*** - *blue*). The ‘+’ marks the individual neuron showing significant baseline change (as assessed by *Two-sample t-test* with *p-value<0.05*). The numbers in parentheses are the percentage of neurons showing baseline modulations in each neuron clusters. The line plot on the top shows the distributions of these changes, and the overall mean with 95% confident interval shown on the top-left of the panel. Except for the paradigm-**N, AM** task (***b***), the remaining task variants (***a, c-d***) display similar trends: neurons from both ***Trans*** and ***Sus(-)*** clusters often displayed an increase of baseline activity in the active state, while no systemic changes of baseline firing rate were observed in ***Sus(+)*** neurons. *, **, ***, *ns* correspond to p-values < 0.05, 0.01, 0.001 and >0.05 of paired *t-test* (overlaid barplots above the barplots) or one sample t-test for non-zero mean distribution (above the barplots), respectively.

**Supplementary Figure 7.**
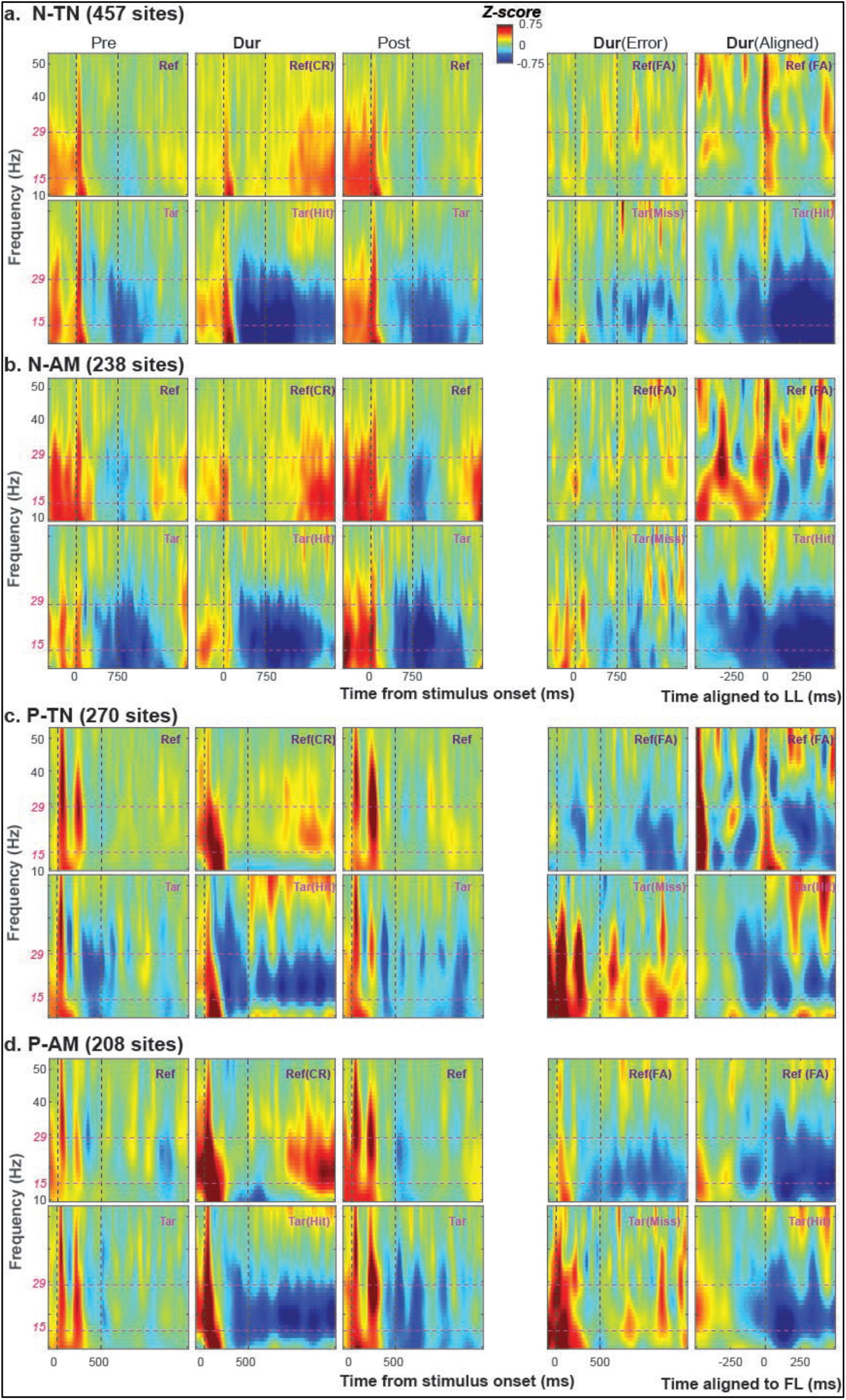
β-band rhythms for difference task variants The power spectrograms extracted from the ***LFPs*** from different task variants are illustrated in plots ***a-d***, corresponding to ***N-TN*** (***a***), ***N-AM*** (***b***), ***P-TN*** (***c***) and ***P-AM*** (***d***) tasks, respectively. The panels (from left to right), display the averaged power spectrogram induced by **Reference** (*top*) and **Target** (*bottom*) stimulus categories during **(i)** *pre-passive*, **(ii)** correct trials in active task engagement *(**CR**/**Hit***) and **(iii)** *post-passive,* and also **(iv)** *the error trials* (***FA****/**Miss***) *in active task engagement* and **(v)** aligned behavioral response trials (*FA* from Reference and ***Hit*** form Target). Note that the β-band power induced by ***FA*** trials shows similar trends to the ***Hit*** trials in P-paradigm (***c, d***), but not the N-paradigm (***a, b***). While the β-band power induced by ***Miss*** trials shows similar trends to Hit trials in N-paradigm, but not in P-paradigm.

## References

Alexander, G.E., Delong, M.R., and Strick, P.L. (1986). Parallel organization of functionally segregated circuits linking basal ganglia and cortex. Ann. Rev. Neurosci. 9: 357–381.

Alexander, G.E., and Crutcher, M.D. (1990). Functional architecture of basal ganglia circuits – neural substrates of parallel processing. Trends in Neuroscience 13(7): 266–271.

Baladron, J. and Hamker, F.H. (2020). Habit learning in hierarchical cortex-basal ganglia loops. European Journal of Neuroscience 52: 4613–4638.

Baladron, J. and Hamker, F.H. (2024). Re-thinking the organization of Cortico-Basal Ganglia-Thalamo-Cortical loops. Cognitive Computation 16:2405–2410.

Banerjee, A., Parente, G., Teutsch, J., Lewis, C., Volgt F.F., and Helmchen, F. (2020). Value-guided remapping of sensory cortex by lateral orbitofrontal cortex. Nature 585: 245–250.

Bariselli, S., Fobbs, W.C., Creed, M.C., Kravitz, A.V. (2018). A competitive model for striatal action selection. Brain Research 1713: 70–79.

Bimbard, C., Demene, C., Girard, C., Radtke-Schuller, S., Shamma, S.A., Tanter, M., Boubenec, Y. (2018). Multi-scale mapping along the auditory hierarchy using high-resolution functional Ultrasound in the awake ferret. Elife 7:e35028.

Bolkan, S.S., Stone, I.R., Pinto, L., Ashwood, Z.C., Garcia, J.M.I., Herman, A.L., Singh, P., Bandi, A., Cox, J., Zimmerman, C.A., Cho, J.R., Engelhard, B., Pillow, J.W., and Witten, I. (2022). Opponent control of behavior by dorsomedial striate pathways depends on task demands and internal state. Nature Neuroscience 25: 345–357.

Bornhoft, K.N., Prohofsky, J., O’Neal, T.J., Wolff, A.R., Saunders, B.T. (2025). Striatal dopamine represents valence on dynamic regional scales. J. Neuroscience: e1551242025.

Bruni, S.B., Giorgetti, V., Bonini, L., and Fogassi, L. (2015). Processing and integration of contextual information in monkey ventrolateral prefrontal neurons during selection and execution of goal-directed manipulative actions. Journal of Neuroscience 35(34): 11877–11890.

Caras, M.L. and Sanes, D.H. (2017). Top-down modulation of sensory cortex gates perceptual learning. Proceedings of the National Academy of Sciences 114: 9972–9977.

Coley, A.A., Padilla-Coreano N., Patel, R., and Tye K.M. (2021). Valence processing in the PFC: Reconcilling circuit-level and systems-level views. International review of Neurobiology 158: 171–212.

Cui, G. et al. (2013). Concurrent activation of Striatal Direct and Indirect pathways during action initiation. Nature 494: 238–242.

David, S.V., Fritz, J.B., Shamma, S.A. (2012). Task reward structure shapes rapid receptive field plasticity in auditory cortex. Proceedings of the National Academy of Sciences, 109: 2144–2149.

DeLong, M.R. (1990). Primate models of movement disorders of basal ganglia origin. Trends in Neuroscience 13: 281–285.

Diesburg, D.A., Greenlee J. D.W., Wessel, J.R. (2021). Cortico-subcortical beta burst dynamics underlying movement cancellation in human. eLife: e70270.

Duan, L.Y., Horst, N.K., Cranmore, A.A.W., Horiguchi, N., Cardinal, R.N., Roberts, A.C., Robbins, T.W. (2021). Controlling one’s world: identification of sub-regions of primate PFC underlying goal-directed behavior. Neuron 109: 2485–2498.

Elgueda, D., Duque, D., Radtke-Schuller, S., Yin, P., David, S.V., Shamma, S.A., and Fritz, J.B. (2019). State-dependent encoding of sound and behavioral meaning in a tertiary region of the ferret auditory cortex. Nat. Neurosci. 22, 447–459.

Engel, A.K., and Fries, P. (2010). Beta-band oscillations – signaling the status quo? Current Opinion in Neurobiology 20: 156–165.

Frank, M.J. (2005). Dynamic dopamine modulation in the basal ganglia: a neurocomputational account of cognitive deficits in medicated and non-medicated Parkinsonism. Journa of Cognitive Neuroscience 17: 51–72.

Friedman, N.P., and Robbins, T.W. (2021). The role of prefrontal cortex in cognitive control and executive function. Neuropsychopharmacology 47: 72–89.

Fritz, J., Shamma, S., Elhilali, M., and Klein, D. (2003). Rapid task-related plasticity of spectrotemporal receptive fields in primary auditory cortex. Nat. Neurosci. 6, 1216–1223.

Fritz, J.B., Elhilali, M., and Shamma, S.A. (2005). Differential dynamic plasticity of A1 receptive fields multiple spectral tasks. J Neuroscience 25(33): 7623–7635.

Fritz, J.B., David, S.V., Radtke-Schuller, S., Yin, P., and Shamma, S.A. (2010). Adaptive, behaviorally gated, persistent encoding of task-relevant auditory information in ferret frontal cortex. Nat. Neurosci. 13, 1011–1019.

Gregorlou, G.G., Rossi, A.F., Ungerleider, L.G., and Desimone, Robert. (2014). Lesions of prefrontal cortex reduce attentional modulation of neuronal responses and synchrony in V4. Nature Neuroscience 17(7): 1003–1011.

Gruber, A.J., and McDonald, R.J. (2012). Context emotion and the strategic pursuit of goals – interactions among multiple brain systems controlling motivated behavior. Frontiers in Behavioral Neuroscience, 6: Article 50.

Hernandez-Martin, E., Kasiri, M., MacLean, J., Olaya, J., Liker, M., Chu, J., and Sanger, T.D. (2023). Globus pallidus internus activity increases during voluntary movement in children with dystonia. iScience 26: 107066.

Hervault, M., and Wessel, J.R. (2025). Common and unique neurophysiological processes that support the stopping and revising of actions. J Neuroscience, 45(13):e1537242025.

Horst, J.t., Boillot, M., Cohen, M.X., and Englitz, B. (2024). Decreased beta power and OFC-STN phase synchronization during stopping in freely behaving rats. J Neuroscience 44(32): e0463242024.

Jahanshahi, M., Obeso, I., Baunez, C., Alegre, M., and Krack, P. (2015). Parkinson’s disease, the subthalamic nucleus, inhibition, and impulsivity. Movement disorder 30(2): 128–140.

Jagadisan, U.K., and Gandhi, N.J. (2022). Population temporal structure supplements the rate code during sensorimotor transformations. Current Biology, 32: 1010–1025.

Koechlin, E., Ody, C., Kouneiher, F. (2003). The architecture of cognitive control in the human prefrontal cortex. Science 302: 1181–1185.

Koketsu, D., Chiken, S., Hisatsune, T., Miyachi, S., Nambu, S. (2021). Elimination of the cortico-subthalamic Hyperdirect pathway induces motor hyperactivity in mice. Journal of Neuroscience 41: 5502–5510.

Komura, Y., Tamura, R., Uwano, T., Nishijo, H., Kaga, K., Ono, T. (2001). Retrospective and prospective coding for predicted reward in the sensory thalamus. Nature 412: 546–549.

Kravitz, A.V., Freeze, B.S., Parker, P.R.L., Kay, K., Thwin, M.T., Deisseroth, K., and Kreitzer, A.C. (2010). Regulation of parkinsonian motor behaviors by optogenetic control of basal ganglia circuitry. Nature 466: 622–626.

Li, B., Nguyen, P., Ma, C., Dan, Y. (2020). Inhibition of impulsive action by projection-defined prefrontal pyramidal neurons. Proceedings of the National Academy of Sciences 117, 17278–17287.

Lisman, J. (2014). Two-phase model of the basal ganglia: implications for discontinuous control of the motor system. Philosophical Transactions of the Royal Society B, 369: 20130489.

Liu, Y., Xin, Y., Xu, N. (2021). A cortical circuit mechanism for structural knowledge-based flexible sensorimotor decision-making. Neuron 109: 2009–2024.

Macedo-Lima, M., Hamlette, L.S., and Caras, M.L. (2024). Orbitofrontal cortex modulates auditory cortical sensitivity and sound perception in Mongolian gerbils. Current Biology 34: 3354–3366.

Meyer, H.C. and Bucci, D. J. (2016). Neural and behavioral mechanisms of proactive and reactive inhibition (Review). Learning and Memory 23: 504–514.

Meyer T., Qi, X-L, Constantinidis C. (2007). Persistent discharges in the prefrontal cortex of monkeys naïve to working memory task. Cerebral Cortex: 17 Supplement 1: i170–176.

Miller, E.K., Lundqvist, M., and Bastos, A.M. (2018). Working memory 2.0. Neuron 100: 463–475.

Mink, J.W. (1996). The basal ganglia – focused selection and inhibition of competing motor programs. Progress in Neurobiology 50: 381–425.

Mittelstadt, J.K, and Kanold, P.O. (2023). Orbitofrontal cortex conveys stimulus and task information to the auditory cortex. Current Biology 33: 4160–4173.

Monosov, I.E. and Hikosaka, O. (2012). Regionally distinct processing of rewards and punishments by the primate ventromedial prefrontal cortex. Journal of Neuroscience 32:10318–10320.

Nambu, A., Tokuno, H., Takada, M. (2002). Functional significance of the cortico-pallidal ‘hyperdirect’ pathway. Neuroscience Research 43: 111–117.

Nambu, A., Chiken, S., Sano, H., Hatanaka, N., Obeso, J.A. (2023). Dynamic activity model of movement disorders: the fundamental role of the hyperdirect pathway. Movement Disorders 38: 2145–2150.

Nelson, A.B., and Kreitzer, A.C. (2014). Reassessing models of basal ganglia functions and dysfunction. Annu. Rev. Neurosci. 37: 117–135.

Nolan, S.O., Melugin, P.R., Erickson, K.R., Adams, W.R., Farahbakhsh, Z.Z., Mcgonigle, C.E., Kwon, M.H., Costa, V.D., Hackett, T.A., Carlson, A.C.C., Constantinidis, C., Lapish, C.C., Grant, K.A., Siciliano, C.A. (2025). Recurrent activity propagates through labile ensembles in macaque dorsolateral prefrontal cortex microcircuits. Current Biology 35: 431–443.

Nonomura, S., Nishizawa, K., Sakai, Y., Nambu, A., Isomura, Y., Kimura, M. (2018). Monitoring and updating of action selection for goal-directed behavior through the striatal Direct and Indirect pathways. Neuron 99:1302–1314.

O’Reilly, R.C., and Frank, M.J. (2006). Making working memory work: a computational model of learning in prefrontal cortex and basal ganglia. Neural Computation 18: 283–328.

Pasquereau, B. and Turner, R.S. (2017). A selective role for ventromedial subthalamic nucleus in inhibitory control. eLife 6: e31627.

Pasquereau, B. and Turner, R.S. (2023). Neural dynamics underlying self-control in the primate subthalamic nucleus. eLife 12: e83971.

Picazio, S., Veniero, D., Ponzo, V., Caltagirone, C., Gross, J., Thut, G., and Koch, G. (2014). Prefrontal control over motor cortex cycles at beta frequency during movement inhibition. Current Biology 24: 2940–2945.

Polyakova, Z., Hatanaka, N., Chiken, S., Nambu, A. (2024). Subthalamic activity for execution and cancellation in monkeys. J. Neuroscience 44: 1911–1922.

Radtke-Schuller, S. (2018). Cyto- and Myeloarchitectural Brain Atlas of the Ferret (Springer International).

Redinbaugh, M.J., and Saalmann, Y.B. (2024). Contribution of basal ganglia circuits to perception attention and consciousness. Journal of Cognitive Neuroscience, 36(8): 1620–1642.

Schiller, B., Gianotti, .L.R.R., Nash, K., and Knoch, D. (2014). Individual differences in inhibitory control – relationship between baseline activation in lateral PFC and an electrophysiological index of response inhibition. Cerebral Cortex 24: 2430–2435.

Schmidt, R., and Berke, J.D. (2017). A Pause-then-Cancel model of stopping-evidence from basal ganglia neurophysiology. Phil. Trans. R. Soc. B 372: 20160202.

Schmidt, R., Leventhal, D.K., Mallet, N., Chen, F., and Berke, J.D. (2013). Canceling actions involving a race between basal ganglia pathways. Nature Neuroscience 16(8): 1118–1124.

Schneider, D.M., Sundararajan, J., Mooney, R. (2018). A cortical filter that learns to suppress the acoustic consequences of movement. Nature 561: 391–395.

Sheikhattar, A., Miran, S., Liu, J., Fritz, J.B., Shamma, S.A., Knaold P.O., and Babadi, B. (2018). Extracting neuronal functional network dynamics via adaptive granger causality analysis. PNAS 115(17): e3869–e3878.

Shin, H., Law, R., Tsutsui, S., Moore, C.I., Jones, S.R. (2017). The rate of transient beta frequency events predicts behavior across tasks and species. eLife 6: e29086.

Sippy, T., Lapray, D., Crochet, S., Petersen, C.C.H. (2015). Cell-Type-Specific sensorimotor processing in striatal projection neurons during goal-directed behavior. Neuron 88: 298–305.

Tatz, J.R., Mather, A., Wessel, J.R. (2023). Beta-bursts over frontal cortex track the surprise of unexpected events in auditory, visual and tactile modalities. Journal of Cognitive Neuroscience 35:485–508.

Vaidya, A.R. and Badre, D. (2022). Abstract task representations for inference and control. Trends in Cognitive Sciences 26:484–498.

Varin, C., Cornil, A., Houtteman, D., Bonnavion, P. & de d’Exaerde A. K. (2023). The respective activation and silencing of striatal direct and indirect pathway neurons support behavior encoding. Nature Communications 14: 4982.

Wessel, J.R., & Aron, R.A. (2017). On the globality of motor suppression: unexpected events and their influence on behavior and cognition. Neuron 93: 259–280.

Wessel, J.R. (2020). B-bursts reveal the trial-to -trial dynamics of movement initiation and cancellation. J neuroscience 40(2): 411–423.

Wessel, J.R. and Anderson, M.C. (2024). Neural mechanisms of domain-general inhibitory control. Trends in Cognitive Sciences 28: 124–143.

Wilhelm, M., Sych, Y., Fomins, A., Warren, J.L.A., Lewis, C., Capdevila, L.S., Boehriger, R., Amadei, E.A., Grewe, B., O’Connor, E.C., Hall, B.J. & Helmchen, F. (2023). Striatum-projecting prefrontal cortex neurons support working memory maintenance. Nature Communications 14: 7016.

Winkowski, D. E., Nagode, D.A., Donaldson, K.J., Yin, P., Shamma, A.A., Fritz, J.B., and Kanold, P.O. (2018). Orbitofrontal cortex neurons respond to sound and activate primary auditory cortex neurons. Cerebral. Cortex 28, 868–879.

Yin, P., Fritz, J.B., and Shamma, S.A. (2014). Rapid spectrotemporal plasticity in primary auditory cortex during behavior. J. Neuroscience 34 (12), 4396–4408.

Yin, P., Shamma, S.A., and Fritz, J.B. (2016). Relative salience of spectral and temporal features in auditory long-term memory. J. Acoustical Society of America. 140, 4046–4060.

Yin, P., Strait, D.L., Radtke-Schuller, S., Fritz, J.B., Shamma, S.A. (2020) Dynamics and hierarchical encoding of non-compact acoustic categories in auditory and frontal cortex. Current Biology 30:1649–1663.e5.

Zikopoulos, B., and Barbas, H. (2007). Circuits for multisensory integration and attentional modulation through the prefrontal cortex and the thalamic reticular nucleus in primates. Rev. Neurosci. 18(6): 417–438.

## Reference

Bagur, S., Averseng, M., Elgueda, D., David, S., Fritz, J., Yin, P., Shamma, A., Boubenec, Y. and Ostojic, S. (2018). Go/No-Go task engagement enhances population representation of target stimuli in primary auditory cortex. Nature Communications 9: 2529.

Foffani, G. & Moxon, K.A. (2004). PSTH-based classification of sensory stimuli using ensembles of single neurons. J Neuroscience Method 135: 107–120.

